# Joint Variant and *De Novo* Mutation Identification on Pedigrees from High-Throughput Sequencing Data

**DOI:** 10.1101/001958

**Authors:** John G. Cleary, Ross Braithwaite, Kurt Gaastra, Brian S. Hilbush, Stuart Inglis, Sean A. Irvine, Alan Jackson, Richard Littin, Sahar Nohzadeh-Malakshah, Minita Shah, Mehul Rathod, David Ware, Len Trigg, Francisco M. De La Vega

**Author notes:** Deceased (John G. Cleary). Current address: Department of Genetics, Stanford University School of Medicine, Stanford, CA 94305, USA.

## Abstract

The analysis of whole-genome or exome sequencing data from trios and pedigrees has being successfully applied to the identification of disease-causing mutations. However, most methods used to identify and genotype genetic variants from next-generation sequencing data ignore the relationships between samples, resulting in significant Mendelian errors, false positives and negatives. Here we present a Bayesian network framework that jointly analyses data from all members of a pedigree simultaneously using Mendelian segregation priors, yet providing the ability to detect *de novo* mutations in offspring, and is scalable to large pedigrees. We evaluated our method by simulations and analysis of WGS data from a 17 individual, 3-generation CEPH pedigree sequenced to 50X average depth. Compared to singleton calling, our family caller produced more high quality variants and eliminated spurious calls as judged by common quality metrics such as Ti/Tv, Het/Hom ratios, and dbSNP/SNP array data concordance. We developed a ground truth dataset to further evaluate our calls by identifying recombination cross-overs in the pedigree and testing variants for consistency with the inferred phasing, and we show that our method significantly outperforms singleton and population variant calling in pedigrees. We identify all previously validated *de novo* mutations in NA12878, concurrent with a 7X precision improvement. Our results show that our method is scalable to large genomics and human disease studies and allows cost optimization by rational sequencing capacity distribution.

## 1. INTRODUCTION

Whole-genome and exome sequencing has been successful in the elucidation of highly penetrant genes in early childhood diseases and is making inroads in complex trait studies entailing thousands of samples Gilisen *et al.*, 2012. Due to its shotgun nature, mis-mapping of short reads in complex genomic regions, and relatively high sequencing error rates, calling variants from human high-throughput sequencing (HTS) data still results in substantial false positives and false negatives (Ajay *et al.*, 2011). The problem is magnified when looking for *de novo* mutations in parent-offspring trios or larger families, as this enriches for sequencing artifacts (Veltman and Brunner, 2012). This is problematic since *de novo* mutations are thought to be responsible for about half of all early neurodevelopmental childhood disorders (Veltman and Brunner, 2012) and likely a similar fraction of neonatal/prenatal cases (Saunders *et al.*, 2012; Talkowski *et al.*, 2012).

Numerous methods have been proposed for the identification of variants from HTS data. Initially, methods were based on frequentist approaches that used heuristic filters (McKernan *et al.*, 2009) but soon were displaced by Bayesian approaches (Garrison and Marth, 2012; DePristo *et al.*, 2011) that leveraged prior information and were able to deal with the uncertainty in the data (Marth *et al.*, 1999). Results from the 1000 Genomes Project provided large-scale empirical validation of these approaches and identified the major sources of artifacts in variant calling: mapping ambiguities due to the complexity of the human genome that result in spurious variants, inflated or non-linear base quality values from the sequencing platforms, library artifacts and sequencing chemistry systematic errors that result in false positives and compositional biases (1000 Genomes Project Consortium *et al.*, 2010; 2012). Nevertheless, due to difficulty of modeling all sources of errors and artifacts variant-calling methods evolved into complex, multistep pipelines addressing separately different problems in the data, including a) mapping and alignment; b) realignment for indels or complex regions; c) base quality recalibration; d) initial variant calling; e) variant quality score recalibration. These methods are complex to use, are slow, and cannot deal effectively with related individuals which are present in family studies and the emerging clinical applications of HTS. In addition, due to the lack of true gold standard samples for evaluating the performance of the diverse variant callers, there is confusion in the field about what caller performs better and under what circumstances, and the true-positive/negative compromise at specific filtering strategies (ORawe *et al.*, 2013).

In order to improve upon these problems, we developed a novel Bayesian network framework which calls variants simultaneously across a pedigree implicitly leveraging shared haplotypes in its members and incorporating a Mendelian segregation model. Here we present how our Bayesian framework escapes combinatorial explosion (as compared to more simplistic approaches), is highly scalable to large pedigrees, can deal with low coverage and missing data, and score *de novo* mutations. In addition, we present a novel approach to identify and genotype locally phased indels and MNPs, simultaneously dealing with spurious variants that may arise in complex regions of the genome. This approach can be applied both to single samples, groups of unrelated individuals, pedigrees, or a combination thereof. Coupled with a fast read mapping and aligning algorithm our approach can process reads-to-variants in a matter of minutes (whole-exome) to a few hours (whole-genome) using commodity hardware and scaling linearly with the number of samples.

To assess our methods and compare to others, we devised a strategy to construct a ground truth data set that leverages the Mendelian segregation of variants and phasing of chromosomal segments from parent to offspring in a large family. With this ground truth and new methods to compare datasets of complex variants which can have different representation, we demonstrate that our approach is not only fast, scalable and accurate, but outperforms other commonly used methods in sensitivity and specificity, and tolerates variations in depth of coverage with minimal detriment to called genotypes.

## 2. METHODS

### 2.1. Mapping and Alignment

Mapping of reads to the reference genome is accomplished by building a 2-bit encoded index of hashes from reads groups and then querying this index using hashes computed on the reference genome. Hashes are constructed from a window *w* of the input sequence with an step or overlap of length *s*. If *S* = *s_0_s_1_*⋯*s_w−1_* is the word to be hashed then the resulting hash is

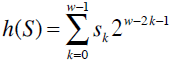

This scheme has the significant advantage of being incremental. Given *h(S)* we can compute

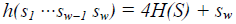

where the multiplication and addition are modulo 2^2w^.

Using this hash value, transformed, sorted set of parallel arrays on sequence data, enables fast numeric search via binary search and other preliminary caching steps. The hash matching is always exact and thus to map a read it is essential that at least one hash generated from the read is found in the index. For the current sequencing platform error rates (about 1% or less), read length (100bp or longer), and polymorphism rate in the human genome, a word size of 20-22 is sufficient to find enough matches for most reads to subsequently trigger an alignment phase in a RAM efficient manner. The step size is set by default to the same as word size, although it could be reduced for additional sensitivity.

#### 2.1.1. Paired-end mapping

The steps for paired-end mapping are:

a. Build an index containing the left/right reads. Filter the index using *repeat-frequency* which removes the hashes that derive from the most highly repetitive locations. By default this is set to 90%.
b. Incrementally analyze each chromosome through a *sliding-window* process exceeding the size of the expected mate-pair insert size, performing read lookup on the first *w* nucleotides. By constraining the search and mating to be the size of the sliding window we guarantee to find all the mated pairs while minimizing the number of spurious locations that reads are aligned against. Each match against the index is then aligned using a multi-stage hierarchical aligner (see below). The alignments that meet alignment quality criteria are output as mated pairs.
c. Once the mated processing is complete, mapping is performed on the remaining (unmated) tags. During searching, we retain the best *top-n* possible locations according to the number of hash hits the read receives at that location. These best locations are then aligned and evaluated against alignment criteria. We use *n* = *5* by default as a good trade-off between memory usage and accuracy.
d. Once mated and unmated mapping is complete, the unmapped reads are output including meta-data regarding the reasons why they are were not aligned.

#### 2.1.2. Multistage hierarchical alignment

Sequence alignment is implemented by a multi-stage hierarchical alignment pipeline. The hierarchical aligner applies incrementally more complex techniques as needed while effectively guaranteeing optimal alignment. The process is:

a. First attempt to perform a perfect alignment, using string comparisons;
b. Then use a substitution only alignment algorithm;
c. Then estimate a lower bound of the sequence difference to identify whether to apply more complex alignment stages;
d. Then apply a seeded aligner (Brown et al., 2004);
e. When required fill in gaps between seeds using a floating point Gotoh improvement to the Needleman-Wunsch algorithm (Gotoh, 1982);
f. If no acceptable alignment is found, leave as unmapped.

The alignment of step e) above is performed with a banded alignment approach to reduce computation time. The default settings allow to aligning indels up to 25bp within 100bp reads but can be set to obtain up to 50bp length indels by adjusting the aligner band width scaling factor.

During mapping, the information for empirical base quality are computed by taking every position in an aligned read, computing covariate variables for the position, and accumulating counts of the number of base matches, mismatches, inserts and deletions associated with those covariate variables. In principle there are many different covariates that could be employed, but by default these covariates are simply reported base quality and read group ID. These calibration tables are written to a file alongside the alignment files. The output of mapping and alignment is a compressed BAM file which includes all uniquely and non-uniquely mapped reads. All recoded mapping locations include an appropriate MAPQ score (H. Li *et al.*, 2008) and is provided in a BAM file (H. Li *et al.*, 2009).

### 2.2. Haploid and diploid variant calling

In what follows we will use the following random variables for a particular locus:

*T* – reference nucleotide;
*S* – read set, set of nucleotides mapped to the locus;
*R* – read nucleotide, a single nucleotide mapped to the locus;
*H* – haplotype – a single nucleotide variant;
*G* – genotype, the one (haploid) or two (diploid) haplotypes of a variant;
*N* – *de novo*, true iff a genotype is possible only as a result of a *de novo* mutation.

What we want to do is to determine the probability of the genotype *G* given the set of nucleotides mapped to a locus and other ancillary information such as the quality scores and variant probabilities. That is, we want to compute *P*(*G*|*S*). Using Bayes formula this can be computed as follows:

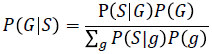

This requires that we compute the conditional probability *P*(*S*|*G*) and the prior *P*(*G*).

Each of the reads in *S* can be considered to be conditionally independent each other. That is, if we know the value of the genotype *G*, then the errors in the different reads are independent of each other. This allows us to decompose the conditional probability distribution (CPD) *P*(*s*|*G*),

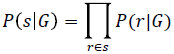

That is, the conditional probability of the whole set of reads is obtained by taking the product of the conditional probabilities of the individual reads.

This now requires computation of the probability of a particular haplotype (say the nucleotide T) given that we have seen a particular nucleotide in the mapped read (say C). If sequencing was error free then this probability would be zero, the only possible explanation would be the nucleotide actually seen in the read. However, there is certainly a non-zero probability of an error. An estimate of this error rate is taken from the quality score supplied in the original sequencing files (e.g. FastQ files). If calibration information is available then this probability will be adjusted by this calibration information (see above). Let the probability of an error be *∈* and consider the case where haploid genotypes are being called then:

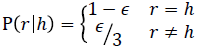

The expression 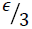 comes from the fact that *∈* is the overall probability of an error and there are three nucleotides other than the correct one each of which is considered equally likely. Such haploid calling is used for the non-PAR regions of the sex chromosomes in males.

For diploid regions of the genome the Bayesian calculation is only slightly more complex. Let the genotype *g* consist of two haplotypes 〈ℎ_1_,ℎ_2_〉 then

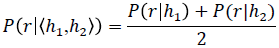

That is the conditional probability is the average of the probabilities for the two constituent haplotypes of the genotype.

#### 2.2.1. Priors

As well as the conditional probabilities described above we also use the priors for the different possible variants, *P*(*G*). This is done using a CPD of transition probabilities *P*(*G*|*t*) that is, *P*(*G*) = *P*(*G*|*t*). The table is generated two ways. The first uses values derived from the results for humans (Levy *et al.*, 2007). The second way of supplying the priors is via a VCF file for a called population. The priors are then computed using the counts of the number of occurrences of the alleles at each location. This calculation has to be careful not to assign a zero probability to any predicted alternative.

Consider the simple case when we are estimating prior probabilities for haploid genotypes. Let *n_h_* be the total number of times allele ℎ has been observed at this locus and 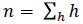 be the total number of observations. Then *P*(*G*) is crudely estimated by

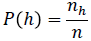

However when *n_h_* = 0 then P(*ℎ*) = 0 which is impermissible. A common way of dealing with this is to add a Laplace correction (Koller and Friedman, 2009):

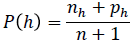

where *p_h_* ≥ 0 and Σ*_h_ p_h._* = 0. In practice this is done by using the default values from above to generate the *p_h_*, that is

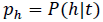

For diploid genotypes Hardy-Weinberg equilibrium is assumed and the diploid priors are computed from the frequencies of the haploid genotypes. The crude estimates are given by

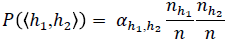

where

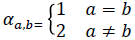

But as above this has a problem when either *n*_*h*_1__ = 0 or *n*_*h*_2__ = 0 An analogous Laplace correction is given by

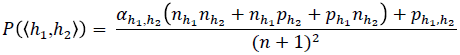

where *p_h_* = *P*(*h*|*t*) and 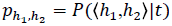.

#### 2.2.2. Mapping errors

The mapper provides an estimate of the probability that a mapping is incorrect in the form of the MAPQ score for each read (H. Li et al., 2008). Let *η* be the probability that the read is mapped incorrectly as indicated by the MAPQ score then we can compute a new CPD P′(*r*|*h*) in terms of the original P(*r*|*h*):

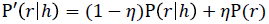

where P(*r*) = Σ*_h_*P(*r*|*h*)*P*(*h*)

#### 2.2.3. Scores

The variant caller outputs its results in a VCF file. The GQ genotype field is provided and follows the standard definition for VCF files. The actual calculated value using the Bayesian probabilities for the genotype is the integer rounded value of

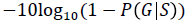

The QUAL info field provides the standard value for VCF files. The actual calculated value using the Bayesian probabilities for the genotype is the integer rounded value of

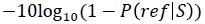

when a non-reference call is made and

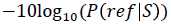

when a reference call is made (normally such a reference call is not placed into the VCF file, however, there are a number of options which can result in some or all such calls being output). *ref* is the appropriate genotype when there is no variation from the reference.

### 2.3. Complex variant caller

It sometimes happens that the alignments achieved during mapping are complicated and disagree with one another in a region of the genome. This can occur when there are indels and other complex variants such as MNPs or a cluster of SNPs in close proximity. In these circumstances while reads may be mapped to roughly the correct location, the details of the alignments may be incorrect.

As a result of this observation we developed a haplotype aware caller. This aims to achieve one or two consistent haplotypes that explain all the reads that overlap a “complex region”. The caller proceeds in three steps:

a. determining (small) complex regions that are candidates for the haplotype caller
b. extracting hypotheses from the alignments for potential haplotypes
c. using a dynamic programming alignment technique that scores the different hypotheses
d. presenting the resulting calls as simple SNPs wherever possible

This can be seen as being intermediate between normal variant calling which believes the mappings given to it and a local assembly technique where all the reads are mapped against each other and a consensus sequence extracted from them.

#### 2.3.1. Complex regions

When determining what regions should be subject to complex calling two classes of loci are determined by the initial SNP calling. First are loci where SNP calls are made or where a low confidence reference call has been made. These are referred to as *interesting* loci. Second are loci where more than one indel has been mapped. These are referred to as *indel* loci.

Indel loci are always treated as complex regions. The complex regions are then grown by merging adjacent regions and interesting loci. An interesting or indel locus is added to the previous region if it is “sufficiently close”. The length for two regions being sufficiently close is composed of the sum of the following terms: i) a constant which defaults to 4; ii) the length of any simple repeats in the gap between the region and the new locus.

The length of the simple repeats is computed by looking for any patterns of 1, 2 or 3 nucleotides that repeat more than once. Finding the repeats may look outside the gap between the existing region and the new locus but the length returned will refer only to the nucleotides inside the gap. Supplementary Table 1 gives examples of this repeat finding calculation.

Once a complex region has been determined, a set of hypothetical haplotypes are extracted from all reads that completely bridge the complex region as determined by the mapped alignment. In some cases the complex regions grow so long that no reads can bridge them. In this case all calling is suppressed in this region.

#### 2.3.2. Alignment of complex regions

Once these haplotypes are extracted the Bayesian calculations are exactly the same as the simple single nucleotide genotypes discussed earlier. The only question that remains is how to compute the term P(*r*|ℎ) that is the probability of a read, *r*, given a haplotype ℎ. This is done by realigning each read against all the hypotheses. The result of this “alignment” is not an alignment *per se* but a posterior probability P(*r*|*ℎ*). The calculation is done using dynamic programming techniques as illustrated in Supplementary Figure 3. First the complex region is replaced by the hypothesis ℎ. The read is then aligned against the reference including this hypothesis. The alignment is done over all possible paths in a parallelogram determined by the start of the original alignment (including an allowance for the start position being incorrect). That is the alignment is against both the hypothesis and the nucleotides surrounding the complex region.

Let the nucleotides in the read be *r_i_* for *i* from 1 to *n*. Let the template with the hypothesis ℎ be written as *t*(ℎ) and the nucleotides in it be *t*(ℎ)*_j_* where *j* has a suitable range. The dynamic programming algorithm incrementally computes the probabilities *p_i_*_,*j*_ defined by

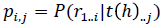

that is, the probability of the first part of the read up to *i* given the part of the reference to the left of *j*. The equation used to incrementally compute *p_i_*_,*j*_ is

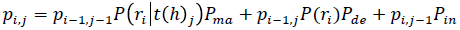

*P*(*r_i_*|*t*(*h*)*_j_*) is a CPD providing the probability of the nucleotide *r_i_* given the reference nucleotide *t*(*h*)*_j_* These probabilities are generated from the probability of a variation occurring in the genome (Levy *et al.*, 2007) and the probability of an error in a read (taken from the calibration information). *P*(*r_i_*) is the probability of the nucleotide *r_i_* taken to be ¼.

The probabilities *P_ma_*, *P_de_*, and *P_in_* are the probabilities of a match, deletion, and insertion respectively (from the rates in the genome and in the calibration files; The match probability is the probability that no insertion or deletion will take place not that there will be a correct match). Note that they sum to 1:

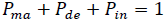

The probability of the read given the hypotheses is calculated by summing over the probabilities along the bottom edge of the parallelogram:

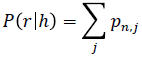

Note that no actual alignments are ever extracted from this, just the sum of the probabilities of all the different paths. This allows considerable speed-up and simplification of the code.

Boundary conditions are applied to limit the size of the parallelogram that the calculation is done over. To the right and left of this the *p_i_*_,*j*_ are set to 0. The vertical height of the parallelogram is given by the read length and the width is ±*w* positions from the start position. *w* currently defaults to 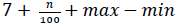 where *n* is the length of the read, *max* is the length of the longest hypothesis, and *min* is the length of the shortest hypothesis. The initial values of the *2w* + 1 positions at the top edge, *p*_0,*j*_, are all set to the same value, ½_*w* + 1_.

As usual during nucleotide alignments the algorithm makes a distinction between starting an indel and extending an indel. To handle this three states (match, insert, and delete) are provided at each position. The probability of another insert (delete) out of the insert (delete) state is higher than the probability of an insert (delete) from the normal match state. This reflects the usual convention that an indel open penalty is larger than an indel extend penalty (Brown, et al., 2004). Supplementary Figure 3 shows the various states and the transitions between them.

#### 2.3.3. Output of complex caller results

After calling has been completed using the haplotype aware caller the output is adjusted to simplify it. For example, sometimes the result is that the reference only should be called, that is, the variant alignments from the mapper were incorrect. In this case nothing is output in the VCF file (although a note is made in a BED file that this was a complex region). If there are common sequences equal to the reference at the ends of the one or two called hypotheses then these are trimmed. If after this it is possible to split the hypotheses into a small number of independent SNPs then this is done.

### 2.4. Adaptive Variant Rescoring

The Bayesian theory for calling and scoring variants assumes a model about how variants occur in genomes and how the sequencing and mapping processes occur. These assumptions are violated in a number of poorly understood ways. For example, there are some reads which are chimaeras of actual nucleotide sequences or are from unassembled parts of a reference genome. To make allowance for these effects a scoring system based on machine learning is used to rescore each variant. The intent is to get the scores to rank the variants in order of their reliability. Then it is possible to filter them on different cut-offs so that subsets of the data with different trade-offs of accuracy and sensitivity can be made.

Our current Adaptive Variant Rescoring (AVR) models are based on Breiman’s random forest ensemble learning technique (Breiman, 2001). We are given a set of training instances over attributes. Each training instance is either positive or negative as determined by external information such as pre-existing baseline. The attributes used are info and format fields appearing in the VCF record together with additional attributes derivable from other attributes. Both real and discrete attributes are supported. The model comprises of a set of decision trees where each tree is trained on a bootstrap sample of the training instances. In building a tree, each node considers some small subset of attributes and a split point is chosen which minimizes the error of the tree with respect to the training set and according to an information gain measure. Training continues recursively on both subtrees until all the instances in the subtree are in the same class, or a predefined minimum number of instances is reached, or there is no further information gain for any potential split. The default models comprise 75 trees with each leaf node representing a minimum of 10 instances. During building 1 + log_2_ (*A*) attributes are considered at each node, where *A* is the total number of attributes in the model. A global weighting factor is applied so that the total weight of positive instances is as close as possible to the total weight of negative instances.

In general it is impossible to provide training sets that are known to be correct. Instead the technique used is to as far as possible enrich the positive examples with true instances and the negative examples with false instances. Because this process is fallible the probability predictions of the models will not converge to the correct probabilities. However, in practice the models provide a good technique for ranking the variants calls.

We built three models that can be used in different situations. The first uses calls from whole genome sequencing of the samples NA12878, NA12891, NA12892 called using our family caller. The second uses calls in the capture regions of exome data for the same samples. Both these models use all of the attributes listed in Supplementary Table 2. The third model uses only the three attributes XRX, GQD, ZY and is intended as a low quality generic model which can be used on most data. One set of training instances are created by taking the supplied calls and checking them against the variant calls published by the 1000 Genomes Project. Calls that agree with one of the samples are treated as positive instances and ones that disagree as negative instances. A second set of training instances are created by checking against the sites published by the HapMap project. Any calls that are at sites included in the HapMap set are treated as positive instances and those that do not as negative instances. Note that it is quite possible for a call to be treated as positive in one set and as negative in another.

During prediction, each instance is presented to each tree. For an instance with no missing values, there is a unique leaf node in the tree corresponding to that instance. The score of the instance is the ratio of the positive instances to total instances at that node as calculated during training. Missing values are handled by constructing weighted sums of both subtrees below the split point corresponding to the missing value. The final AVR score is the average of the scores returned by the individual trees.

### 2.5. Joint Bayesian calling in pedigrees

Given the genotypes *G_f_*, *G_m_* and *G*_1_ and the sets of mapped reads *S_f_*, *S_m_* and *S*_1_ for the father, mother and one offspring respectively the joint distribution for the combination is:

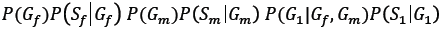

Later it will be clearer if we write this expression for the joint distribution somewhat more abstractly as

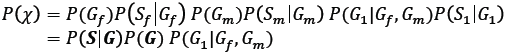

*P*(*χ*) is shorthand for the joint distribution over all variables, and ***S*** and ***G*** indicate the random variables for all sets of reads and all genotypes respectively (Figure 1).

**Figure 1.**
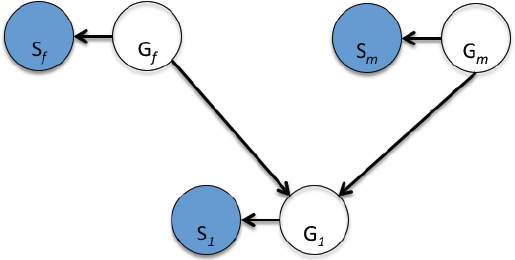
Bayesian network for a parent-offspring trio.

Using Bayesian rules it is possible to extract from this expression the posterior distributions that we are looking for, in this case, *P*(*G_f_*|***s***), *P*(*G_m_*|***s***) and *P*(*G*_1_|***s***) the posterior distributions respectively for the father’s, mother’s and offspring’s genotypes given all the mappings of reads.

All of the factors in the joint distribution except one are provided by the single sample calling. *P*(*G_f_*) and P(*G_m_*) are the prior probabilities for the genotypes. These are computed from tables of known nucleotide frequencies and if a priors VCF file is supplied, from the allele counts there. The factors *P*(*S_i_*|*G_i_*) link the nucleotides seen in the mapped reads to the genotypes. What is new here is the factor *P*(*G*_1_|*G_f_*, *G_m_*) which is a CPD linking the genotypes of the parents to the offspring’s. This factor takes into account Mendelian inheritance, mutation and the type of chromosome, that is, whether it is an autosome or a sex chromosome. Supplementary Table 3 shows part of the CPD for SNPs in an autosome. Similar tables are used for the sex chromosomes depending on the sex of the offspring. The three cases are: Y chromosome in a son which is inherited only from the father; X chromosome in a son which is inherited only from one of the mother’s two alleles; and the X chromosome of a daughter which is inherited from the father’s only X and one of the mother’s two X chromosomes. The details of these inheritance patterns are specified as part of the configuration of the reference sequence for a genome and can be configured for other patterns in non-mammalian species.

#### 2.5.1. de novo mutation scoring

In the strictly Mendelian inheritance of Supplementary Table 3, an instance such as *P*(〈A,T〉|〈A,C〉, 〈A,C〉) will have a probability of 0. However, allowance is made for a *de novo* mutation between the parents and the offspring with a probability of 10^−9^. In order to ensure that the sums of the probabilities in the CPD are still 1, the other cases all have a slightly reduced probability. This small probability means that significant evidence is needed from the mapped reads before such a *de novo* call will be made.

Whenever a call is made which can only occur as a result of a *de novo* mutation a special DNP score is calculated. This is the posterior score *P*(*N*_1_|***S***) where *N*_1_ is a true false value indicating that the offspring’s genotype can only occur as a result of a *de novo* mutation and ***S*** is all the variables for the parents’ and offspring’s sets of reads. As usual in VCF files, it is output as a Phred score. The value of *N*_1_ is computed deterministically from the genotypes ***G*** so the deterministic CPD *P*(*N*_1_|***G***) can be included in the joint distribution without disturbing the other variables

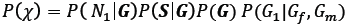

*P*(*N*_1_|***S***) is then computed in the normal way by summing over all the values of ***G*** and applying Bayes theorem.

The Bayesian analysis above can be extended easily to families with more offspring. The joint distribution then becomes:

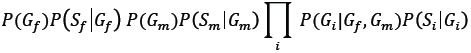

that is, the factors for all the offspring are multiplied in. In strictly theoretical terms there is no problem with this formulation, however, if the calculations are implemented naïvely the execution time will increase exponentially with the number of offspring. However, the offspring are conditionally independent of each other given the parents genotypes, which permits the calculations to be done in time *O*(*n*^3^) where *n* is the number of potential genotypes for a single sample.

### 2.6. Multi-generational pedigrees

Pedigrees can be much more complex than a nuclear family with two parents and some number of offspring. Supplementary Figure 1 shows an extended pedigree and Figure 2 the corresponding Bayesian network.

**Figure 2.**
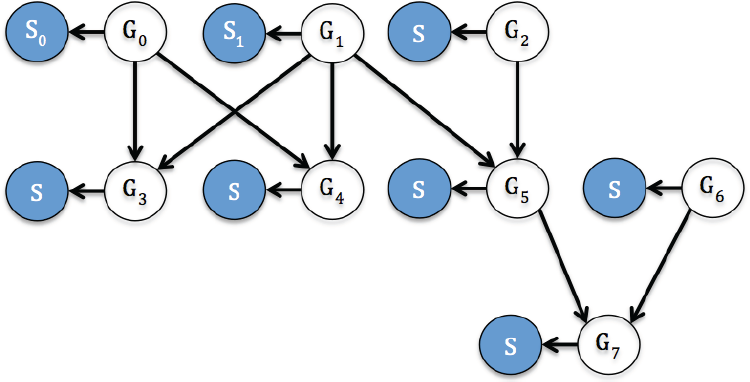
Bayesian network for an extended pedigree.

To express the joint distribution for such a pedigree it is necessary to introduce some notation. Let *i*^↑^ be the (unique) father of sample *i* and *i*^†^ be the (unique) mother of sample *i*. *i* is a root if it has no father or mother. It is assumed that if one parent is present then the other is also. This can be achieved by adding a sample which contains no reads (*S_i_* = ∅) if sequencing data from such a sample is absent.

The factor for a root node contains the prior for the genotype and the CPD for the reads given the genotype, *P*(*s_i_*|*Gi*)*P*(*G_i_*). The factor for a non-root node contains the CPD for the genotype given the genotype of the parents and the CPD for the reads given the genotype, 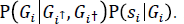 All of these factors have been met already in the nuclear family case.

Combining all these factors the joint distribution is

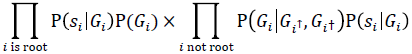

As before there is a danger that a naïve calculation of the posterior distributions *P*(*G_i_* | ***s***) will explode exponentially with the size of the pedigree. If the pedigree contains no inbreeding then standard techniques for variable elimination in Bayesian networks are guaranteed to converge to the required posterior. For example, using a message passing algorithm (Shafer and Shenoy, 1990) the computation can be arranged as two passes. First a backward pass proceeds from the leaves towards the roots. It incrementally computes a partial joint distribution that includes all the samples that are descended from each node (this calculation for a parent requires only the result of the backward pass for each of the offspring). Second a forward pass from the root nodes to the leaves calculates a partial joint distribution that includes each sample and all samples that are not descended from it (this calculation requires the result of the forward pass for each offspring as well as the backward pass for each of its siblings and half-siblings). Finally the results of the forward and backward pass for each sample *i* are combined to compute *P*(*G_i_* | ***s***). Practical techniques for implementations are discussed in (Koller and Friedman, 2009) and (Pearl, 1988).

If there is some inbreeding then this algorithm breaks down (there will come a point where it is not possible to proceed with the forward pass of the algorithm). This can be solved at the cost of further computation by iterating the backward and forward passes. Whenever a result is required in a pass that hasn’t already been computed, then an old value from a previous pass (or a default prior) is used. The iterations continue until there are small changes in the calls and their scores.

### 2.7. Joint Population Calling

So far we have focused on the additional information that can be gained by knowing the relationships of the samples. Even in the case when there is no known familial relationship extra information can be gained by knowing that other individuals have a particular variant. This is done by using the frequency that a genotype occurs at a locus in the whole population as an estimate for the prior when doing Bayesian calling, that is *P*(*G_f_*) or *P*(*G_m_*) in a single family.

The way this is computed can be viewed as a form of the Expectation Maximization (EM) algorithm. First all the samples are called (at one locus) using default priors and any family information that is available. That is, the isolated samples are called as if they are single samples and the families and larger pedigrees use the inheritance information. Once all the calls in the different samples are done the number of times each allele occurs is counted. For haploid calls these are used to give a frequency estimate for each allele with a suitable Laplace correction to deal with cases never seen in the data. For diploid calls a similar procedure is used where the estimated frequency for a particular pair is proportional to the product of the counts for the two alleles. The Laplace corrections in this case are as described previously.

Once the new estimates have been made the calling is repeated. This EM cycle of calling and estimation is repeated until the calls do not change. In practice this usually takes only 2 or 3 iterations although there are a small number of cases in large populations that are stopped by a cut-off of 25 iterations.

It should be emphasized that the population calling is using both the pedigree information where it is available and an assumption of Hardy-Weinberg equilibrium through the complete population including both the family founders and isolated members. In cases where there is both inbreeding and the EM algorithm is being used, then the iterations required by the EM algorithm and the Forward-Backward algorithm are combined.

If external priors from a previous calling are provided, then the priors estimation at each iteration includes the counts both from the population under consideration and from the external source. Thus it is possible to incrementally build up calls for a large population by using the results from previous calling runs to improve the results from another run.

### 2.8. Complex variant calling in joint sample analysis

Haplotype aware calling is also used in the joint caller. The main differences are that all samples can contribute potential haplotypes and that the different haplotypes are evaluated using the same Bayesian logic as used for SNVs.

Because there can be very large numbers of samples there can potentially be large numbers of putative haplotypes. The haplotypes are pruned to include at most 6 alternatives ranked on how often they occur. In some exceptional circumstances there may be many haplotypes with the same counts and it may not be possible to easily get a set of 6, in these cases no attempt is made to call the complex region.

### 2.9. Benchmarking

#### 2.9.1. Simulations

To evaluate our method we simulated a WGS dataset for a parent-offspring trio by constructing the four haplotypes of the founders from the Human reference sequence (hg19) and applying polymorphism using as a template the data from the 1000 Genomes Project to a rate of 10^−6^. We then recombined the parents haplotypes with a crossover rate of one in 100Mb, ensuring that at least one crossover per chromosome arm existed. From this template we simulated 2 × 100bp paired-end reads with an average insert size of 350bp and a sequencing error rate similar to that of the Illumina platform with a higher probability of error at the 3’-end of the reads and an average error rate of 1%. Reads were simulated randomly across the genome to and average depth of 5X.

#### 2.9.2. Gold standard data

In addition to simulations, we analyzed a set of “gold standard” WGS data from a 3-generation CEPH/Utah family of 17 members (Supplementary Figure 4) produced by Illumina Inc. (San Diego, CA) as part of their “Platinum Genomes” resource (http://www.illumina.com/platinumgenomes/). DNA from lymphoblastoid cell lines obtained from the Coriell Institute were sequenced with the HiSeq^®^ 2000 system to 50X average depth using 2 × 100bp libraries of ∼350bp insert size. Raw data was downloaded from the European Nucleotide Archive (Acc. Nos. ERA172924 and ERA185981). We aligned reads and performed calls in 3 nuclear family subsets and the entire pedigree for comparison (see Supplementary Figure 4).

#### 2.9.3. HTS Data Analysis

We analyzed simulated or real HTS data with a multithreaded Java implementation of our methods that is part of the commercially available rtgVariant v1.2 software (Real Time Genomics, Inc., San Bruno, CA). Software was run on commodity dual-quad Intel servers running the Linux operating system (Amazon EC2, m3.2xlarge instances running CENTOS, Intel Xeon E5-2670 eight core, 34 GB RAM, launched by StarCluster). Reads were mapped in groups of up to 50 million reads jobs using eight cores, to the Human genome reference hg19 with decoys as described by the 1000 Genomes Project (1000 Genomes Project Consortium *et al.*, 2012). BAM files were merged as needed and passed to the variant caller providing, when necessary, a PED file describing the pedigree relationships and sex of the sample. Variant calling was parallelized by splitting jobs by chromosomes and each job was multithreaded in eight cores. As described in the results, samples were either called independently or in groups. The output VCFs were either analyzed in its entirety, or after filtering with an AVR score of ≥ 0.15 for whole-genome data model, or ≥ 0.45 for whole-exome data model. Compressed VCF files with the different calls sets produced for this study are available at the NCBI trace archive at: ftp://ftp-trace.ncbi.nih.gov/giab/ftp/data/NA12878/variant_calls/RTG/.

For comparison purposes we also aligned the gold-standard data with BWA v0.6.1 and called variants with two other methods: GATK Unified Genotyper v1.7 (DePristo *et al.*, 2011), and Samtools-hybrid v0.1.7 (H. Li *et al.*, 2009). These tools where used with their default parameters, including filtering down to the first tranche of VQSR for GATK. In addition, to compare with another approach for family-aware variant calling, we reanalyzed the VCFs from Samtools-hybrid with PolyMutt v0.16 (B. Li *et al.*, 2012).

#### 2.9.4. Orthogonal reference data

A set of reference data for comparisons was obtained for the sample NA12878. These include:

a. Illumina OMNI v2.5 SNP Arrays data from the 1000 Genomes Project (ftp://ftp.1000genomes.ebi.ac.uk/vol1/ftp/technical/working/20120131_omni_genotypes_and_in tensities/
b. CDC Get-RM high quality variants downloaded on May 27th 2013 (Ball et al., 2012) (ftp://ftp.ncbi.nlm.nih.gov/variation/get-rm/current/NA12878_high_quality_variant.vcf.gz)
c. NIST Genome-in-a-Bottle arbitration datatset v 2.15 (Zook et al., 2013) (ftp://ftp-trace.ncbi.nih.gov/giab/ftp/data/NA12878/variant_calls/NIST)
d. Variant calls for CEPH/Utah pedigree obtained by Complete Genomics Inc. using their sequencing-by-ligation chemistry (Drmanac et al., 2009). (ftp://ftp2.completegenomics.com/).
e. Validated *de novo* mutations in sample NA12878, both germline and cell line somatic from (Conrad et al., 2011) and (Ramu et al., 2013).

#### 2.9.5. Identification of cross-overs and phasing of large pedigree

Given access to data from a large pedigree, such as the CEPH/Utah pedigree with 11 offspring, it should be possible to deduce the relative phasing of both the children and the parents using phase by transmission. Such phasing can then be used to construct a ground truth dataset towards which one can test the quality of variant calls made by different callers.

The task of establishing the phasing begins with a set of diploid calls of both the parents and all 11 offspring filtered to high quality for all variant and reference calls. The basic assumption here is that recombination cross-overs are infrequent and that there is a small probability that individual calls are incorrect. Calls which are non-Mendelian, where both parents are homozygous, or where all calls (parents and children) are heterozygous, are discarded as likely artifacts.

For each autosomal sequence and for the X-chromosome, a greedy algorithm was used to establish blocks of consistent phasing. It was assumed that these blocks would be broken by erroneous calls or by cross-over events, these are dealt with later. The phasing for each child can be described in terms both of which parent each half of a diploid genotype was taken from and of a haplotype within the parents (given the available information the haplotypes in the parents cannot be assigned to the grandparents and so are labeled arbitrarily). The haplotypes are labeled *f_a_*, *f_b_*, *m_a_* and *m_b_*, two for the father and two for the mother (the more complex situation in the X-chromosome is explained below). Thus a phasing for a child will be an assignment of the alleles in its genotype to an ordered pair such as *f_a_*, *m_a_* or *m_b_*, *f_a_*. Genotypes are output in the notation of the VCF file format, that is, 0 for the reference allele and 1, 2, 3 etc. for alternative alleles. Given the calls at one locus, it is seldom possible to uniquely determine the phasing. For example, if the father’s genotype is 0/0 and the mother’s is 0/1, then it is impossible to place any constraints on the father’s phasing although a constraint is possible on the mother’s. Table 1 below gives all the possible cases that can occur (given that one of the parents must be heterozygous and other cases can be generated by switching the parents and/or switching the labels within the parents genotypes). Table 1 shows the two situations mentioned above when the parents are 0/0 and 0/1. If a child has the genotype 0/0, then there are two possible phasing labels: *f_a_/m_a_* or *f_b_/m_a_*. A similar situation arises if the child has a 0/1 genotype. If the parents are both heterozygous with a 0/1 and 0/1 genotypes then the situation is more complex. If the child has a 0/0 (1/1) genotype then there is only a single possible labeling *f_a_/m_a_*(*f_b_/m_b_*). However, if it is heterozygous then there are two labelings, *f_a_/m_b_* and *f_b_/m_a_*, which partially constrain both the possibilities for the father and mother. The remaining cases where there are more than two alleles all allow of only a single labeling.

**Table 1.** Phasing labels given parent and child genotypes.

| Parents |  | Children |  |  |  |
| --- | --- | --- | --- | --- | --- |
| $f_a/f_b$ | $m_a/m_b$ | | | | |
| 0/0 | 0/1 | 0/0 | 0/1 |  |  |
| | | $f_a/m_a$ | $f_a/m_b$ | | |
| | | $f_b/m_a$ | $f_b/m_b$ | | |
| 0/1 | 0/1 | 0/0 | 0/1 | 1/1 |  |
| | | $f_a/m_a$ | $f_b/m_a$ | $f_b/m_b$ | |
| | | | $f_a/m_b$ | | |
| 0/0 | 1/2 | 0/1 | 0/2 |  |  |
| | | $f_a/m_a$ | $f_a/m_b$ | | |
| | | $f_b/m_a$ | $f_b/m_b$ | | |
| 0/1 | 1/2 | 0/1 | 0/2 | 1/1 | 1/2 |
| | | $f_a/m_a$ | $f_a/m_b$ | $f_b/m_a$ | $f_b/m_b$ |
| 0/1 | 2/3 | 0/2 | 0/3 | 1/2 | 1/3 |
| | | $f_a/m_a$ | $f_a/m_b$ | $f_b/m_a$ | $f_b/m_b$ |

The situation for the X chromosome is simpler. The father is homozygous so there are only two non-trivial cases as shown in Table 2. The pseudo autosomal regions (PAR) on the X and Y chromosomes are dealt appropriately as diploid although no attempt was made to bridge the phasing in the non-autosomal regions with that in the PAR regions.

**Table 2.**
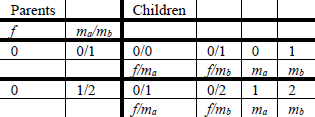
Phasing labels for X chromosome.

| Parents |  | Children |  |  |  |
| --- | --- | --- | --- | --- | --- |
| $f$ | $m_a/m_b$ | | | | |
| 0 | 0/1 | 0/0 | 0/1 | 0 | 1 |
| | | $f/m_a$ | $f/m_b$ | $m_a$ | $m_b$ |
| 0 | 1/2 | 0/1 | 0/2 | 1 | 2 |
| | | $f/m_a$ | $f/m_b$ | $m_a$ | $m_b$ |

We started with high quality genotype data for the WGS of the pedigree, by calling NA12878 jointly with her parents as a trio, similarly for the other parent, sample NA12877, and concurrently calling the 11 offspring, but pointing the latter to “dummy” parents with no data to avoid biases due to joint Mendelian calling of the parents and the offspring. Genotype data was filtered by AVR 0.15> across all samples, including reference calls. Our method greedily constructs blocks of compatible loci moving 5’(p) to 3’(q) through a chromosome. A block contains a set of labelings for each child. A new block is started by adding the labelings for the next locus as described above. For subsequent *loci* an attempt is made to intersect the labelings for the children with that of the block so far. To do this it may be necessary to flip the labelings for the parents (there are four ways this can be done flipping *a* and *b* for each of the mother and father). There are two possible results for this: If a way can be found to make the next *locus* compatible with the current block then the labeling on the block can be updated, often to a less ambiguous labeling, and the process continues. If it is not possible to find a compatible labeling then the current block is recorded and a new block is started with the labeling at the next *locus*. A new block will be forced this way whenever there is a cross-over or when an error has occurred in the calling.

Once a sequence of such blocks have been established, a dynamic programming algorithm is used to find an optimal path through the blocks. This can treat each block in four ways, each of which has an associated cost. If the next block is compatible with the previous (non-error) block included in the path then it is added to the path with no cost. If the next block is incompatible then three things might be done: i) Mark the block as in error, that is, the calls in the block are incorrect and the block does not contribute to the phasing. The cost is equal to the number of variant loci in the block. ii) Start a new path with this block; the cost is 1,000. This may be necessary if there has been a region which is hard to map or where there have been multiple cross overs between variant loci. iii) If it is possible mark the new block as being a cross over; the cost is 100. This is only possible if the last (non-error) block in the path is compatible with all the children except one which has a single phase difference in one of the father or mother.

As is usual in dynamic programing the optimal path is retrieved after a forward search pass by choosing the lowest cost path and following back links. Also, if two paths converge to the same state (the same phase labeling at the same locus) then the higher cost one can be forgotten. Once this process is finished, we have identified a set of intervals that include candidate recombination cross-overs (see Supplementary Files 1-5 for an a graphic representation of the crossover across chromosomes of the offspring coming from for each parent and the path contiguity for each chromosome). A histogram of the number of cross-overs split by chromosome and parent is shown in Supplementary Figure 5 and shows that as expected, larger chromosomes and the mother present more cross-overs (Kong *et al.*, 2010), which supports our results.

#### 2.9.6. Comparing Variant Call Files

When evaluating how correct a variant caller is, we assume that we have three pieces of information: a reference sequence (for example an assembled genome); a baseline set of variations on the reference; and a called set of variations on the reference. The called sequence will be the best possible one if it correctly includes everything in the baseline (the true positives), and has no incorrect calls (false positives) and no calls it has missed (false negatives). However, a complication arises due to possible differences in representation for indels and MNPs in two different call sets. This often happen for indels within repeats, differences in how many bases are included in MNP calls, and how the variant are aligned in respect to the 5’ or 3’ ends of the reference.

In order to deal with these problems, we devised a method that “replays” the variants from both the baseline and called set of variants to a given reference genome assembly. Once the variants are replayed, we then search for the best path that maximizes true positives and minimize false positives and false negatives. A path through a call sequence is a selection of subset of calls. The idea is that the calls included as correct in the baseline and called paths will be equal (after being replayed). The calls excluded from the baseline will correspond to false negatives and the calls excluded from the called sequence will be classified as false positives. The calls in the baseline path and the called path (which always agree) are classified as true positives. The method searches through all possible paths in both the baseline and called sequences and finds the pair of paths that maximize true positives and minimize errors. Potentially there are an exponential number of paths to be explored, however if the alternative (replayed) paths converge at the same position on the reference, then the one which minimizes the number of errors up to that point can kept and the others discarded. In practice this happens frequently keeping the memory and processing requirements reasonable. Note that our method considers ploidy when comparing variants, and thus to match, variants should have the same genotype. All heterozygous variants are treated as non phased and evaluated accordingly.

When comparing variant calls the number of true positives plus the number of false negatives should equal the total number of calls in the baseline. Generally, each called variant will have a corresponding baseline variant but due to the representation differences mentioned above, there can be a 1 to many relationships between baseline and called mutation. To keep number of true positives plus the number of false negatives equal to the total number of calls in the baseline, each called true positive call must be weighted. To avoid other pitfalls due to ambiguity when looking for equivalences in repeat regions, we perform weighting within “sync points”. A sync point is a location where diploid baseline and called path are at the same position on the reference and they are not currently in the middle of any variant location. An optimization in the best path creation skips all the genomic locations which does not contain any variants, thus the sync points occurs just before the next available variant. Once all the sync points are created, each called mutation is weighted using following formula:

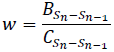

where *B* is the number of baseline variants between the current (*S_n_*) and previous sync points (*S_n−1_*) and *C* is the number of called variants between the current and previous sync points. For examples of the best path searches, variant matching, and weighting, see Supplementary Note 1. This method is implemented in the vcfeval software, which is freely available as part of the rtgTools package (downloadable at ftp://ftp-trace.ncbi.nih.gov/giab/ftp/tools/RTG/).

#### 2.9.7. Evaluating calls vs. pedigree phasing labels

Once the phasing has been established in a large pedigree as described above, any set of calls can be checked against the phasing labels. Given an input VCF file the checker produces a new annotated VCF file. Four things can happen at each variant locus:

a. If the locus falls in a region which has not been phased then it remains unchanged except for the addition of a “PHO” filter;
b. If the calls do not fit the phasing pattern then the following are added: i) a “PHI” filter; ii) an INFO field also called “PHI” which contains a comma separated list of characters for each child, “I” for a child inconsistent given the genotypes of the parents and “C” for consistent; iii) and an INFO field “PHIC” which contains the count of the number of children which are inconsistent;
c. If the calls do not fit the phasing pattern but they will if just one sample has its call changed are treated specially (see below for details);
d. If the call does correctly phase then each of the GT fields is phased and an INFO flag “PHC” is added and a “PHQ” INFO field.

Loci which are phase inconsistent but where it is possible to arrive at a consistent phasing by changing the call for one of the parents or children are treated specially. These may be the results of random minor errors and can be rescued by giving the site the benefit of the doubt. Two lines are generated in the VCF file at the same locus:

a. The first line has information about the repaired call: i) a “PHR” INFO field with the name of the sample that was repaired and the original unchanged GT; ii) a “PHQ” INFO field; iii) the GT field of the repaired sample is changed to the new repaired call
b. The second line has information about the original unrepaired calls: i) a “PHI” INFO field as described above; ii) a “PHI” and a “PHR” (PHased Repair) filter, all other information remains unchanged from the original.

It is possible for a region to not have been phased for two reasons. At the ends of chromosomes the blocks on the lowest cost path may have been marked as an error. This leaves the regions at the end without a reliable phasing. If the first or last block is included in the lowest cost path then it is assumed that the phasing reaches all the way to the end of the chromosome. A second similar way that this can occur is when it has been necessary to break the low cost path because it has not been possible to explain the blocks with a cross-over. This can occur because two cross-overs occur so close to each other that they cannot be resolved by the call set used to construct the phasing (unlikely), or there is a region where there are many errors and it has not been possible to find a path through it. Each cross-over will occur between two of the original loci used to construct the phasing. So, for a locus which is being checked and which lies between the two original positions, it is ambiguous which of two possible phasings should be used. Both are checked and if either of them is correct then the locus is marked as being correctly phased.

The phasing of the children’s GT fields follow the usual convention by putting the father’s allele to the left of a “|” and the mothers allele to the right, e.g. “1|0”. The phasing for the parents use phase groups (because the grand parents have not been included in the analysis it is not known which grandparent the alleles came from). The phase groups are restarted whenever there is a break in the lowest cost path through the blocks.

An annotated VCF with the phasing-consistent variants from the Illumina Platinum data called with rtgVariant, called the Segregation Phasing (SP) standard, is available at the NIST GiaB repository at: ftp://ftp-trace.ncbi.nih.gov/giab/ftp/data/NA12878/variant_calls/RTG/.

### 2.10. Joint calling at different depths of coverage

In order to experimentally test whether joint variant calling in pedigrees could compensate differences in depth of coverage and thus enable a more cost-effective sequence capacity distribution, we undertake a design where the parents are sequenced at half the usual coverage, whereas the proband (offspring) is sequenced at high depth. DNA from lymphoblastoid cell lines of the individuals obtained from the Coriell Institute was sequenced with the HiSeq^®^ 2000 system either to ≈100X or 50X average depths using 2 × 100bp libraries of ∼350bp insert size by Edge BioSystems (Gaithersburg, MD). For the full depth data, four samples were indexed and enriched together with the Agilent^®^ SureSelect All Exome Kit v 4, and loaded on a single HiSeq lane. In the case of half depth coverage, 8 samples where combined and placed together on a lane for sequencing (samples were enriched in sets of 4).

We aligned reads and performed calling using the following configurations:

a. Full Coverage (FC) Family Calling - Parents and child were sequenced to ∼100X each and using pedigree information for variant calling
b. Reduced Coverage (RC) Family Calling - Parents were sequenced to ∼50X each and child to ∼100X and pedigree information was used for variant calling.
c. Reduced coverage for all (ARC) – all members of the trio sequenced to ∼50X.
d. Singleton Calling - FC or RC Parents and child data was called independently, without knowledge of their relationship.

Raw variants were filtered to those on the target regions of the enrichment assay and with the exome-recommended AVR cut-off of 0.45 (Illumina exome model).

## 3. RESULTS

### 3.1. Simulations

For an initial evaluation of our method we simulated HTS paired-end reads of 100bp length with an average insert size of 350 bp, to an average depth of 5X for a parent-offspring trio. Figure 3 shows a receiver operator characteristic (ROC) curve showing a comparison between calling the offspring independently (singleton), jointly with her parents using the population calling without considering the family relationships, and jointly but using the family calling, sorted by GQ score. These results demonstrate that the family caller performs much better that the other two cases having a greater area under the curve (AUC) and being able to retrieve more true positives at the expense of fewer false positives at any given GQ score threshold. This improvement is particularly notable at reduced depth of coverage vs high depth coverage (data not shown).

**Figure 3.**
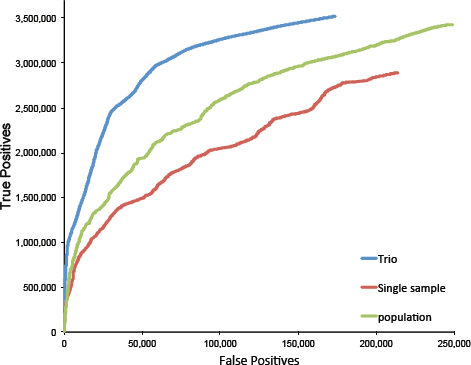
ROC curve of call sets from a simulated parent-offspring trio. Data is for the offspring called as either a single sample (offspring; red line), with its parents using the population caller (green line), or jointly as a trio with the family caller (blue line).

### 3.2. Analysis of gold standard samples

To better asses the performance of our method with real HTS data and its associated artifacts, we analyzed the data from a set of “gold-standard” samples from a widely used CEPH/Utah pedigree sequenced with 2 × 100bp PE Illumina reads to about 50X (see Methods; Supplementary Table 4). We focus our analysis on NA12878, a female in the second generation, for which extensive orthogonal validation data exists including fosmid-end Sanger sequence data (Kidd *et al.*, 2008), Complete Genomics WGS data, Illumina OMNI SNP-array genotype data (1000 Genomes Project Consortium *et al.*, 2012) and experimentally validated germline and cell-line somatic *de novo* mutation data (Conrad *et al.*, 2011; Ramu *et al.*, 2013). This sample is also part of an effort to develop reference materials for HTS laboratory validation, and a high-confidence set of variants constructed through an arbitration process from data of multiple sequencing platforms, is available (Zook *et al.*, 2013). Table 3 shows that, compared to singleton calling, the family caller produced more SNV/indel/MNP calls while maintaining good quality, as judged by commonly used quality metrics such as Ti/Tv, Het/Hom ratios, and dbSNP array concordance. Note that the data was filtered using an AVR ≤ 0.15 cut-off for our caller, whereas for GATK the 1^st^ tranche of VQSR was used, as recommended (DePristo *et al.*, 2011).

**Table 3.**
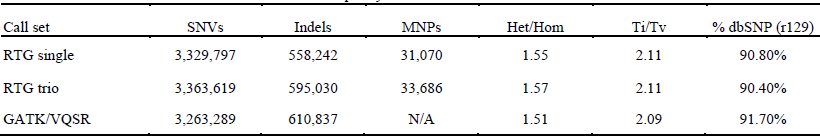
Statistics and quality metrics for different call sets for NA12878

**Table 4.** Concordance with orthogonal datasets for NA12878 call sets.

| Call set | 1000 Genomes<br>Project OMNI Poly<br>(TP%) | 1000 Genomes<br>Project OMNI<br>Mono (FP%) | Get-RM<br>(TP %) | GiaB<br>(TP%) | GiaB-BED<sup>1</sup><br>(TP%) |
| --- | --- | --- | --- | --- | --- |
| RTG single | 97.50% | 0.10% | 97.40% | 89.80% | 93.40% |
| RTG trio | 97.50% | 0.24% | 97.00% | 90.50% | 94.10% |
| GATK/VQSR | 97.80% | 0.17% | 87.80% | 88.40% | 92.50% |

<sup>1</sup>High confidence BED region.

We then compared the calls sets with reference datasets obtained by orthogonal methods for the sample NA12878, including SNP OMNI array data generated by the 1000 Genomes project (1000 Genomes Project OMNI Poly, for polymorphic sites), the Get-RM high confidence calls (Ball *et al.*, 2012), and the NIST Genome-in-a-bottle arbitration calls (Zook *et al.*, 2013). Although all call sets show good concordance with the orthogonal data, the family caller showed slightly better agreement with the arbitration data sets of the NIST, in particular in the high confidence regions (GiaB-BED). As a proxy for false positives we report the variants discovered across ≈50,000 sites in the OMNI SNP array that appear momomorphic in the 1000 Genomes Project samples (DePristo *et al.*, 2011; 1000 Genomes Project Consortium *et al.*, 2012). All numbers are very low, and although the family caller appears to have higher call rate in these sites, manual inspection of a random set of alignments suggest that many of these calls are real, but either small indels or MNPs, which presumably cannot be detected in the microarray platform.

While these comparisons suggest that joint trio calling performs better, they are filtered at the default quality score threshold and don’t reveal the performance across the entire scoring range. We thus constructed ROC curves using as a baseline (or ground truth) either the arbitration dataset developed by the NIST (Figure 4, Panel A) or the independently obtained variants calls produced by Complete Genomics (Figure 4, Panel B). As can be seen in Figure 4, the family caller outperforms the single-sample calling (greater AUC) allowing making a better trade-off between true positives and false positives at realistic thresholds. As compared to GATK-UG, our family caller in general shows greater AUC in the range up to the end of the VQSR 1^st^ tranche, but there is variation depending on the baseline used. Less differences are seen with the NIST GiaB data set; this could be the result of biases to GATK calls and/or Illumina platform data during the construction of this version of the GiaB baseline whereas the CGI data is completely independent.

**Figure 4.**
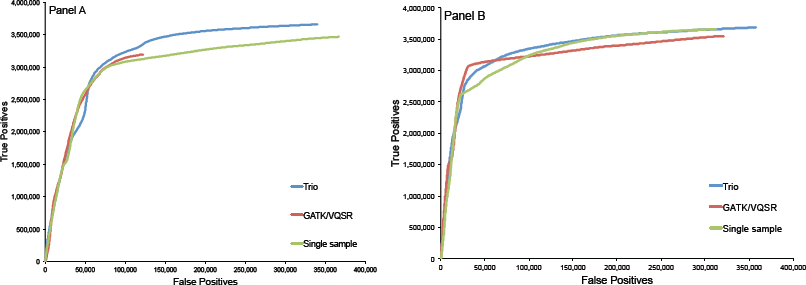
ROC curve of different calls sets for NA12878 vs the CGI calls (Panel A) or NIST GiaB calls (Panel B) as baseline. Calls are for the following methods: Single sample (offspring; green); Trio (blue); and GATK/VQSR single sample (offspring red). Our calls are sorted by AVR score (see Methods), GATK are sorted by VQSLOD score.

### 3.3. Assessment of calls with a pedigree-derived ground truth

In view of the pitfalls seen with the aforementioned baselines, we developed a ground truth call dataset taking advantage of the extensive Mendelian segregation of variants in the last generation of the CEPH/Utah Pedigree to assess and compare the genotype calls from different methods and parameters. The rational is that if a variant is segregated to the next generation following Mendel laws, this suggests that the variant is a true positive. There are many situations where a technical artifact, either due to the sequencing technology or analysis methods, would appear to segregate in a Mendelian fashion to the offspring of a trio. However, barring simple errors that can be filtered out, it is very unlikely that variants that segregate in a Mendelian fashion and within the correct haplotype context across 11 offspring are artifacts.

In fact, it is possible to compute an estimate of the probability that a particular locus will agree with the phasing by chance given the distribution of genotypes in the offspring. Since there are *d* different genotypes across both the parents and children and that the number of times each of these genotypes occurs is *n*_i_ and *n* = Σ*_i_ n_i_* then the probability is

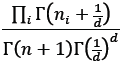

In the case where the phasing is ambiguous the probability is the sum of the probabilities for the two possible phasings. Supplementary Table 4 gives some examples of the probabilities that result from this equation where there are 11 children. It includes the most extreme cases where the genotypes are all “1/1” which results in a probability of 1, and some other cases where there is a third allele and where the probability is as low as 10^−5^.

We then identified cross-overs and phased the entire chromosomal segments of the large CEPH 1463 family as described in the Methods. Once this phased framework was constructed, we took the set of genotype calls from our joint calling of the large family of the CEPH/Utah pedigree and tested for consistency with this phasing (Table 5). Over 99.99% of the calls fall within the phased segments. As can be observed, close to 20% of the raw, unfiltered calls are inconsistent with the phasing framework, but when a threshold of AVR > 0.15 is used for filtering, this drops dramatically to just 0.3%. Some variants may not be phase-consistent due to a random error in single individual and we may want to rescue those, giving the benefit of the doubt to that sample, and repairing the genotype (see Methods). As it turns out, only a few percent of variants can be rescued in this way.

**Table 5.** Phase consistency of variants called on the CEPH 1463 large family with the joint pedigree caller.

| Call Set | Raw |  | AVR >0.15 |  |
| --- | --- | --- | --- | --- |
|  | n | % | n | % |
| Phase consistent | 5,224,138 | 77.35 | 4,606,574 | 99.28 |
| Phase inconsistent | 1,329,189 | 19.68 | 13,951 | 0.30 |
| Repaired | 200,450 | 2.96 | 19,197 | 0.41 |
| Calls inside phased segments | 6,753,777 | 99.99 | 4,639,722 | 99.99 |

For comparison, we analyzed phase consistency of the calls produced by the GATK unified genotyper (v1.7) from BWA alignments (both raw and VQSR 1^st^ tranche filtered), and for the calls produced by Complete Genomics (CGI) for the same pedigree (final calls only; (Drmanac *et al.*, 2009)). Figure 5 shows that the joint caller produces 10X less inconsistent calls (i.e. false positives) as compared with GATK after filtering, while maintaining high sensitivity (cf. Table 3). Surprisingly, the CGI calls exhibited a very high number of inconsistent calls (cf. Supp. Table 5 an 6).

**Figure 5.**
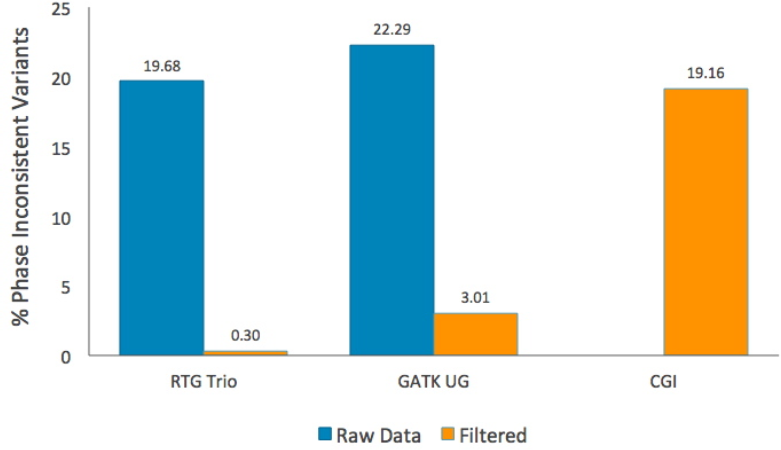
Phase inconsistent calls of the CEPH 1463 large family called by the RTG joint caller, and GATK UG from the Illumina Platinum data, or by the CGI platform and analysis software. Filtered calls for RTG are AVR > 0.15 and for GATK UG is the VQSR 1^st^ tranche.

We also analyzed separately the rate of phase-inconsistent calls (i.e. false positives) for indels and MNPs of different lengths called by our best joint calling call set. Figure 6 includes the results when filtering the calls with the recommended AVR ≥ 0.15 cut-off and shows that although the overall number of FPs is small, the number increases with longer indels, in particular longer insertions. Comparatively, MNPs have more false positives than indels, whereas single-nucleotide variants have a very low false positive rate of ≈ 0.2%.

To construct a ground-truth variant call set using the phasing information, we simply aggregated all phase-consistent calls identified by this process after removing some likely artifacts and calls where all samples were heterozygous (see Methods). Using this ground truth, termed SP (for Segregation Phasing), we evaluated several calling configurations and callers, focusing on the sample NA12878. Figure 7 shows ROC curves for different call sets using this ground truth.

**Figure 6.**
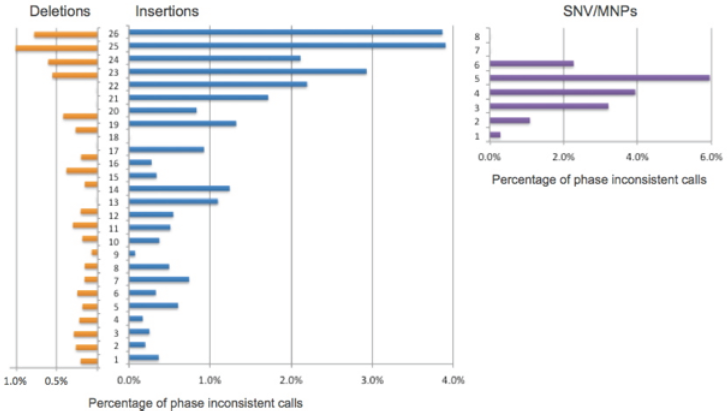
Percentages of phase inconsistent calls (false-positives) for indels and MNPs, segregated by length. Data from NA12878 called as a trio and filtered with AVR ≤ 0.15.

**Figure 7.**
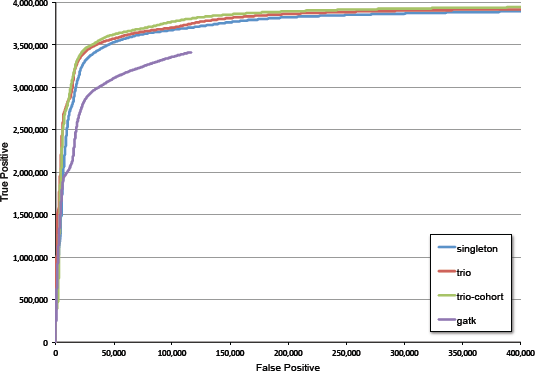
ROC curves of different call sets vs the segregation-phasing baseline. The datasets compared include rtg singleton (blue), rtg trio (red); rtg trio-cohort (green; when the trio is called in the context of the other samples but without relationships), all sorted by AVR; and the 1^st^ VQSR tranche of GATK-UG (v1.7; purple), sorted by VQSLOD score.

We observe an improvement in the AUC when the sample is called with the trio caller as compared with the single sample calling (Figure 7). A further improvement in AUC is achieved when the trio is called in the context of the remaining samples of the pedigree, even when the relationships between the trio and the rest of the samples are not provided (trio-cohort). This improvement is mostly on the total number of variants called, the bulk of the difference with the simple trio curve being on complex calls. We also observe that our method significantly outperforms the GATK-UG single sample calls and it is able to provide much more sensitivity at a fixed false positive rate. For example, at the recommended AVR ≥ 0.15 cut-off, the FP is just 1.9% at a sensitivity of 93.9% (using the trio-cohort curve from Fig. 7), while for the 1^st^ tranche of VQSR the FP is 3.2% at a sensitivity of 87%.

### 3.4. Mendelian errors

One of the major aims of jointly calling pedigrees is the reduction of Mendelian errors. These errors are significant and are problematic in downstream data analysis for many applications. We thus enumerated the genotypes incompatible with Mendelian inheritance across the trio samples. Table 6 shows that the family caller dramatically reduces the number of Mendelian inconsistency errors (MIEs) as compared with single sample approaches. It should be noted that this is accomplished without resorting to filtering and indeed the family caller returns more calls than single-sample calling (cf. Table 3).

**Table 6.** Mendelian errors across trio samples for different callers.

| Calling Method | Mendelian inconsistent calls |  |
| --- | --- | --- |
|  | n | % |
| Singleton caller | 335,626 | 5.62% |
| Family caller | 4,870 | 0.09% |
| GATK/VQSR | 351,904 | 9.2% |
| PolyMutt | 9,283 | 0.32% |

We compared the results of our method with those from PolyMutt (B. Li *et al.*, 2012), a likelihood based method aimed at improving variant calls in a trio-aware fashion and identify *de novo* mutations. PolyMutt requires as input a call set from another variant calling, so in this case we used the output from SamTools-hybrid (H. Li *et al.*, 2009). As can be seen in Table 6, PolyMutt reduces the number of Mendelian errors to 0.32%, but still 3.5-fold more than our method.

### 3.5. Imputation of missing samples

One consequence of the Bayesian approach is that it is possible to impute a genotype for one of the family members even if there is no or very small coverage for it. This is particularly true for the parents when there are multiple offspring, these can strongly constrain the parents genotypes.

Figure 8 shows that as we add more offspring to the computation we can recover the calls from the mother increasingly better, until an imputation of over 96% is achieved when all eleven offspring is used (the other father is always present in the analysis). Further, the total number of false positives decreases dramatically as more information is available from the additional offspring, reaching a reasonable trade-off around 3-4 offspring, and showing a small improvement when all 11 offspring are included. This demonstrates not only the ability of the family caller to leverage information from other family members to improve the calls across all individuals, but also to deal with missing sample data which could be useful in many situations.

**Figure 8.**
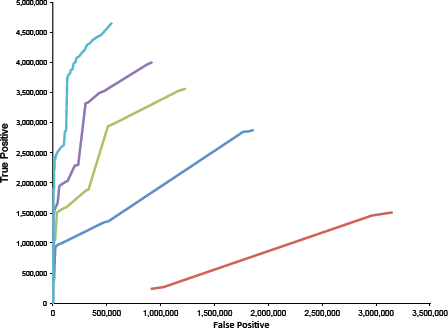
ROC curve for the imputation of calls for NA12878 as a function of the data of her offspring. Each line represents the ROC curve as a number of offspring is included in the computation (red = 1; dark blue = 2; green = 3; purple = 4; light blue = 11). The baseline is the calls of the sample obtained when calling the entire family. The other parent (NA12877) is always present.

### 3.6. Identification of *de novo* mutations

One of the main advantages of the joint family caller is the ability to score putative *de novo* mutations in offspring simultaneously with calling variants. Table 7 shows that without the ability to consider the parents jointly, a large number of heterozygote calls in the offspring whose variant allele appears not to be present in *either* parent is present in the call set. However, we know from prior studies that the number of true *de novo* mutations that appear in the germline should be closer to about 100 (B. Li *et al.*, 2012; Conrad *et al.*, 2011). Even considering that this data is obtained from a cell line where somatic *de novo* mutations exist (about 1000), the *de novo* candidates in single sample call sets are hugely contaminated with false positives. However, the family caller identifies just under 3,000 candidates; a reduction of 60X.

**Table 7.**
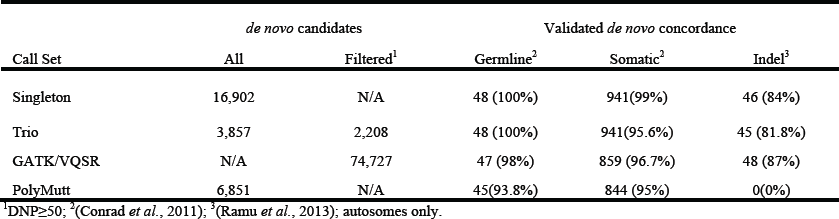
*de novo* mutation identification in NA12878

The 1000 Genomes Project experimentally validated a subset of the *de novo* germline and somatic mutations in the NA12878 trio that we can use as a reference (Ball *et al.*, 2012; Conrad *et al.*, 2011). As compared with the validated *de novo* mutations set, we observed 100% and 99% sensitivity in detecting the reported germline and somatic *de novo* single-nucleotide mutations, respectively (note that the cell line batch may be different and thus may have slightly different somatic mutations). Through the analysis of variant segregation to the third generation, we confirmed 99% of the Conrad *et al* (Zook *et al.*, 2013; Conrad *et al.*, 2011) germline mutations (somatic variants do not segregate, as expected). Importantly, the high *de novo* sensitivity was achieved while reducing the number of candidate *de novo* mutations by greater than 6-fold without using *ad hoc* filters. We derived a specific quality score for *de novo* mutations (DNP, see Methods). Filtering with this score (e.g. DNP ≥ 50) the number of candidate mutations can be reduced further (Supplementary Figure 6). We also analyzed the *de novo* mutations identified by PolyMutt (B. Li *et al.*, 2012) in the same data. As shown in Table 7, PolyMutt identifies twice the number of candidate *de novo* mutations (i.e. half the specificity) and is less sensitive in identifying the 1000 Genomes Project validated *de novos*, as compared to our joint trio calling method. In addition, PolyMutt only identifies single-nucleotide *de novo* mutations.

As we add additional offspring to a family, the possible *de novo* mutations candidates should be more constrained, both by the additional information provided by the additional sample, and the low probability of de novo mutations being shared across siblings. Supplementary Table 7 shows that as we add additional siblings, the number of *de novo* mutations called in NA12885 (a sample from the 3^rd^ generation of the pedigree) decreases as expected.

### 3.7. Multigenerational pedigrees

Expanding the analysis to multigenerational pedigrees requires new methods that avoid combinatorial explosion to be practical. We implemented a belief propagation algorithm in our Bayesian network model that allows propagating priors beyond nuclear families. To evaluate the advantage of this method we simulated data for a multigenerational pedigree at different depths of coverage. Supplementary Figure 6 shows the pedigree structure we used to simulate data. On a first instance all individuals were simulated at a depth of 5X, and in a second example we simulated the grand parents to 30X and the rest to 5X. As shown in Supp. Figure 8, the AUC of the ROC for either the mother and the first father (F1) increases in the call set from the family caller as compared with single sample calls (HTS is simulated to a depth of 5X). To better understand the impact of prior propagation from one generation to another, we simulated 30X coverage for the grandparents and 5X for the rest of the pedigree. In Supp. Figure 9 we observe that while the benefit of family calling for the grandparents is small (they have already high coverage), there is a substantial benefit for the other samples, such as F1 and even M. The impact in distal samples is smaller (data not shown). We also tested our method in the complete CEPH/Utah pedigree data. Supplementary Table 8 shows that calls have good quality very similar to the results for nuclear families, while maintaining a MIE rate of 3.6% in the entire pedigree.

**Figure 9.**
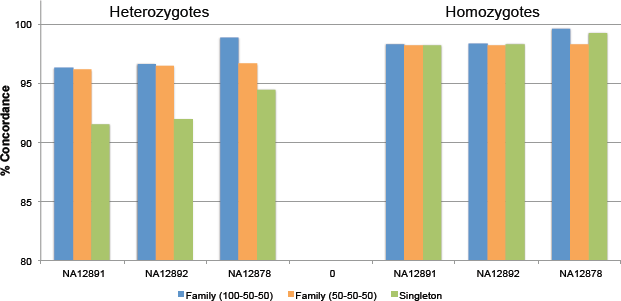
Heterozygote and homozygote recall ratios for the samples of the trio at full vs recued coverage of parents and compared with the single sample calling.

### 3.8. Planning sequencing capacity with joint calling

Joint calling improves the accuracy of called variants by incorporating data across the pedigree members. We hypothesized this can be applied to more cost-effective study designs by distributing sequencing capacity across individuals. One such designs we tested involves cost reduction in trio WES or WGS, a typical clinical application, by reducing the depth of coverage of the parents 50%, thus reducing the cost of the study by one-third as compared to the full coverage.

As compared to family calling on full coverage parents (FC), the reduced coverage WES set (RC) produced equally high quality variant calls, as judged by commonly used quality metrics such as Ti/Tv, Het/Hom ratios, and dbSNP/OMNI array concordance for NA12878. High sensitivity was achieved concomitant with a low 1–2% false positive rate as assessed by variants called at monomorphic sites in the 1000 Genomes Project OMNI SNP-array (Supplementary Table 9). In contrast, when the samples were called as singletons, the call set of the half-coverage parents show a reduction in the number of variants identified and a reduction in sensitivity as measured by discovery of 1000 Genomes Project OMNI Poly and Complete Genomics (CGI) reference SNV sites. In addition, a reduction in the Het/Hom ratio suggests increased undercalled heterozygotes as would be expected due to lower coverage.

Given that the point of including the parents’ information in many settings is to use their genotypes to infer mode of inheritance (e.g. simple recessive, compound heterozygotes) and *de novo* mutations, it is important to demonstrate that genotyping accuracy is preserved in the reduced coverage samples. This is particularly important for heterozygote calls, as they will suffer more from reduced depth. In the following analysis, the full coverage (FC) calls were considered as the baseline and we separated the calls into heterozygous and homozygous groups.

We compare the genotype concordance for reduced coverage (RC) when called jointly, or when the samples are called as singletons. Figure 9 shows that the family caller data shows 96% and 99% heterozygous concordance in the parents and child respectively, recovering from any detrimental effect of the lower coverage. However, in singleton calls, heterozygote concordance drops to 91% in the reduced coverage parents. Note that even in the offspring, which is sequenced to high depth, there is also a drop in heterozygote concordance to 94%, because it is not benefiting from the information from the parents such as in the family calling case. As expected, the homozygote concordance is much less affected in either case.

A question to consider is whether reducing the coverage in the parents of a trio impairs the ability to reduce MIEs in the dataset. To determine this, we counted the Mendelian inconsistency errors on each individual for the full coverage and half coverage study designs. As shown in Supp. Table 10, a very low number of MIEs are observed in either case, showing that reduced depth in the parents does not impact the ability to correct those errors. In contrast, when the offspring have been called independently as a singleton, a high number of errors are observed (∼30-fold vs family). This shows that even at reduced coverage, there is a persistent advantage of using the trio information in variant calling.

Preserving the ability to identify *de novo* mutations under reduced coverage is an important criterion to measure the benefits of this approach. Out of the validated *de novo* mutations set from Conrad *et al.* (Conrad *et al.*, 2011), 14 mutations are present in the exome sequencing assay. We are able to call all of these correctly for both the FC and RC cases. This high sensitivity is accompanied with a 30X reduction in Mendelian errors (see Supp. Table 11).

### 3.9. Performance

Another aim we had when we developed this method was to enable fast analysis of HTS data, from single samples, to trios, to populations, and large multigenerational pedigrees, in a unified fashion. Therefore, we analyzed the performance of our method with commodity hardware to inform on its ability to handle the growing volume of HTS data becoming available. Mapping and alignment was performed as described in Methods for each sample in an average of 13 hrs cumulative wall clock hours/sample on a commodity Linux server. In contrast, mapping with BWA averaged 97 cumulative wall clock hr/sample.

Supplementary Table 10 shows the wall clock times for different variant calling configurations of the CEPH/Utah samples. Our results show that we can call variants from 50X WGS sample in 7.2 hr cumulative wall clock time per sample (or 0.6 hr wall clock with parallelization in a cluster), which is over 65-fold faster than other methods (e.g. GATK-UG v 1.7, data not shown). The compute time is linear with the number of family members for the nuclear families, whereas in the combined family and population calling (3 nuclear families) and extended pedigree calling the iterative process increased the run times to 22 and 38 cumulative wall clock/hr per sample (1.5 & 1.4 wall clock/hr/sample parallelized), respectively, still 10-times faster than other software. It should be noted that our methods do not require sorting of the BAM files (the mapper already sorted the reads by location), de-duplication, or realignment, additional time-consuming steps in most common pipelines and that contributes to file footprint bloating.

## 4. DISCUSSION

The analysis of pedigrees allowed the identification of a large number of disease genes through linkage mapping. As new high-throughput genotyping technologies arrived, the hunt for disease susceptibility variants switched to case/control genome-wide association studies that resulted in a large number of associated variants, but with low effect and frequently not fine mapped to the actual causal variant. As useful those results can be to inform disease pathways, it is becoming apparent that a significant fraction of heritability unexplained by GWAS hits could be caused by rare variants (Manolio *et al.*, 2009). The novel HTS platforms make now affordable to explore such rare variation by whole-genome or exome sequencing, and family structures are now important tools in this research. In addition, WES has been very successful to elucidate causal genes for highly penetrant Mendelian diseases and currently is being deployed in diagnostic procedures of early childhood diseases. Recent reports suggest that *de novo* mutations could account for over 50% of early neurodevelopmental diseases in outbred populations besides recessive and dominant inheritance modes (Veltman and Brunner, 2012). The identification of such *de novo* mutations and in general the analysis of these cases requires or greatly benefits from sequencing and concurrent analysis of the parents.

In spite of this, variant identification and genotyping methods from HTS data often ignore the family relationships between the samples and the fact that the same haplotypes are being redundantly sequenced. This is an important piece of information that can be leveraged to reduce the inevitable errors that occur in HTS variant calling due to the nature of the method (a shotgun), and the many artifacts due to the short length of the reads, the complexity of the human genome, and the systematic biases of the HTS platforms. There have been some attempts to perform “family-aware” variant rescoring for improving the calls sets or specifically the identifications of *de novo* mutations (Ramu *et al.*, 2013; B. Li *et al.*, 2012; Cartwright *et al.*, 2012; Peng *et al.*, 2013). However, these methods are *post-hoc* (reanalyze previously called data), not principled, difficult to use, and slow. In contrast, the method we present here utilizes an extensible Bayesian network framework that permits fast joint family calling leveraging information across related samples using Mendelian inheritance priors, and can also call variants in single samples, groups of unrelated samples, and a combination of pedigrees and unrelated individuals. Our method scores *de novo* mutations, is scalable to multigenerational pedigrees, and includes a novel haplotype-aware algorithm to resolve complex regions that harbor indels, MNPs, or can produce spurious calls. Our results with empirical data from gold standard samples from a CEPH/Utah pedigree demonstrates that the family caller is superior to single sample approaches, producing high quality calls and reducing Mendelian inconsistency errors to very low levels (Tables 3 & 4). This is accomplished without filtering but instead correcting the undercalled heterozygotes in some of the samples using otherwise weak evidence that would have been dismissed. In fact, the family caller returns additional variants, at any given quality threshold, which with the additional information are now rescored higher (Table 3 and Fig. 4).

We show a very high concordance with orthogonal reference data for sample NA12878 suggesting excellent sensitivity and specificity. However, these reference data produced by other methods are not infallible and have pitfalls. For example, the invariant sites from the OMNI genotyping data produced by the 1000 Genomes Project has been widely used as a proxy for false positives. We found a slight increase in OMNI monomorphic sites in the trio calls. When we analyzed alignments by hand for a sample of such variants we find reasonable support for about half of these, which frequently are MNP, or small indel calls, and are unlikely to be typed correctly by the SNP array. To improve upon this, we constructed a ground truth for the CEPH large family by identifying recombination cross-overs on the offspring, extending and connecting inferred haplotype blocks, and then putting back called variants on the haplotype framework. Those variants that were consistent with the haplotype framework, or whose inconsistency can be removed by correcting a single genotype, were assumed to be true positives and added to the ground truth; otherwise they were assumed to be false positives arising from mapping artifacts or systematic sequencing errors. We developed this Segregation Phasing (SP) standard using the rtgVariant calls from the Illumina Platinum WGS data of the pedigree, removing the relationship information of the last generation to avoid biases due to joint Mendelian calling. We were able to reproduce the phasing framework starting from the GATK/VQSR calls from the same samples (data not shown), confirming our results are not biased by our caller. Furthermore, the standard can be augmented by adding phase-consistent calls from multiple callers and sequencing platforms to create a gold standard reference dataset that can used for sequencing pipeline assessment. The SP standard and the methodology developed can thus be a valuable complement and input to other ongoing efforts by the genomics community such as the NIST Genome-in-a-Bottle Project (Zook *et al.*, 2013).

The ROC curves of the different call sets versus the SP baseline (Figure 7) revealed that while the trio calling reduces significantly MIEs and improves the AUC vs single calling, and significantly over the GATK-UG single sample calls, calling other samples simultaneously even if their relationship is removed improves further the AUC. This improvement comes in the form of additional complex calls (indels and MNPs). We explain this as the result of the complex caller using all alignments across the samples to prune the most unlikely complex call hypothesis and highlights the benefits of calling groups of samples whether they are related or not.

We could have completely eliminated Mendelian inconsistent calls, however we allow for a small probability of *de novo* mutations (a prior of 10^−9^, consistent with estimated mutation rates per generation; Conard et al., 2011). This allows capturing true *de novo* variants in offspring while significantly reducing the background of false positive *de novos*. While high sensitivity can be achieved by simply reporting variants that pass less stringent accuracy thresholds (and in so doing increasing substantially the number of MIEs), our family calling achieves high sensitivity concurrently with a 10X reduction in MIEs (MIE; cf. Table 7). Further reduction of false positives can be achieved by filtering with the DNP score, improving precision even further. We evaluated our *de novo* detection using validated mutations from the 1000 Genomes Project. It is noteworthy that these mutations were identified from trio data sequenced to a depth of only 20X, in Phase I of the project. Thus, it is conceivable that additional mutations can be identified with the higher coverage data we examined. Accordingly, we screened our candidates by segregation to the next generation to identify novel germline *de novo* mutations. By removing those candidates that don’t segregate, or segregate only in one or all of the eleven offspring, we rounded up about 200-250 putative novel *de novo* mutations in the NA12878 sample, which await further validation (data not shown). We also identified *de novo* mutations in the 3^rd^ generation offspring, and observe that the number of candidates decreases with the additional information of the siblings as expected (Supplementary Table 7). Comparatively, our method outperforms PolyMutt, another method we tested that aims to identify *de novo* mutations, as reflected in our higher TP and lower FP rates, can detect *de novo* MNPs and indels (cf. Table 7), in addition of being much faster (data not shown).

A possible pitfall of our method is that in regions of copy number variation (CNV) simple Mendelian inheritance is not expected. Indeed the identification of clusters of MIEs in SNP arrays data was used to first identify genome-wide CNVs (McCarroll *et al.*, 2008). Nevertheless, our Bayesian network model includes a factor for copy number of the template. This is normally set to diploid, but it can be changed to accommodate haploid calls in sex chromosomes in males (except for the PAR region in X), and can certainly be changed for regions of CNV if those regions are known *a priori*. In addition, one may want to infer the copy number from the data and indeed our model allows for estimating the CPD of the copy number if desired. Such implementation is aimed for future developments.

Another area of further improvements is in the identification of indels and MNPs (complex variants). Our analysis with the SP standard showed that long indels, in particular insertions, and MNP exhibit higher false positives than small indels and SNVs. Firstly, identifying such variants is confounded by numerous mapping artefacts that occur in real data beyond the paralogy and repeat structures in the human genome. For example, the human genome reference is not complete (misses centromeres, telomeres, and heterochromatin), and the sample we sequence may also have novel insertions, CNVs, and alternative haplotypes (e.g. HLA) This material is present in the reads and are sometimes mapped into close homologues creating spurious variants. Secondly, there is very little empirical validation data for indels and MNPs to serve as ground truth. The 1000 Genomes Project has done some statistical validation of indels, but it was done in pools encompassed only samples not sequenced to full depth (1000 Genomes Project Consortium *et al.*, 2012). There is also some fosmid-end Sanger sequencing data available for NA12878 (Kidd *et al.*, 2008). However, calling indels in this data is not trivial. Finally, indel detection is limited by the initial alignments that, due to mapping reasons, cap the indel alignment length (e.g. due to banding). Currently, our default banding allows initial alignments of up to ∼25 bp, although the complex caller could extend these further during variant calling (see Supplementary Figure 10 for a histogram of the length distribution of detected indels). If additional indel sensitivity is desired, the banding can be increased at the expense of slower alignment time. Alternatively, additional hypotheses for complex variant calling can be obtained by other means, such as local assembly of reads, either those aligned directly during mapping, or those placed in the approximate location due to the mapped position of their mate.

We demonstrate that through the use of joint pedigree calling a significant reduction of sequencing costs can be achieved when analyzing trios by reducing depth of coverage of the parents to half. This is done with little trade-off in accuracy and retaining the benefits of joint family analysis, namely: i) higher sensitivity, ii) a reduction in Mendelian errors, and iii) identification of *de novo* mutations with 60X precision over singleton analysis. In this analysis we focused in the case where only the parents’ sequencing depth has been reduced to not compromise the integrity of the proband data, and because it lend itself better to the experimental workflow used in the sequencing. Other schemes are possible; for example all samples could be reduced equally to, for example, 2/3 of the depth and results are very similar (cf. Supp. Table 9).

The Bayesian network framework we developed is capable to infer other parameters, such as ploidy (CNV) at variant sites, and the model can be naturally expanded to deal with other relationships between samples that are present in different study designs. An instance is the case of cell lineages where one cell type is derived from other but essentially share most of the genomic variants, for example in iPS cell derivation and differentiation, or in gamete differentiation. Another important instance is in the case of somatic mutation identification in cancer tumor analysis, where the primary tumor is derived from the germline tissue and can produce subsequent metastatic tumors. The Bayesian network model can easily accommodate these relationships, incorporate contamination from germline tissue in the tumor samples (cellularity), and changes in ploidy, while jointly analyzing all the data to identify germline and somatic mutations. These are aspects of future work.

Our methods are very efficient and fast: coupled with a hash-based fast mapping and hierarchical alignment method, whole genome sequence data of a parent-offspring trio sequenced to 50X depth (2 × 100 bp) can collectively be analyzed from reads to variant calls in ∼22 hours on a single commodity server, and is amenable to large-scale parallelization for further speed improvements. In summary, our results show that joint pedigree calling outperforms singleton and population variant calling in pedigrees, allows for the identification of *de novo* mutations with greater specificity, and is scalable to large genomics and human disease studies. We believe analytical advances like this are crucial for the adoption of genomes and exomes in large family disease studies and clinical settings.

## Acknowledgements

We would like to thank Chad Harlan (LIC), George Weinstock (Washington University), and Carlos Bustamante (Stanford University) for fruitful discussions. We are grateful for the feedback and encouragement of Jason Blue-Smith, Stephen Lombardi, and Graham Gaylard (RTG), at different phases of this work. We are indebted to Michael Eberle (Illumina, Inc.), and Steve Lincoln (formerly Complete Genomics, Inc.) for releasing publicly the CEPH/Utah pedigree WGS data we utilized and providing related information. We also thank the 1000 Genomes Project Consortium and Justin Zook (NIST) for the reference data used in this work. We dedicate this work to the memory of John G. Cleary.

## Author’s Contributions

JGC, SI, SAI, LT and FMV conceived methods. JGC, RB, KG, SI, SAI, AJ, RL, MR, DW, and LT developed and implemented algorithms. FMV and JGC conceived the validation study and analysis strategy. SNM, RB, BSH, MS, MR, RL, and LT performed data analysis. SNM, MS, RL performed production runs. JGC and FMV wrote the manuscript.

## SUPPLEMENTARY TABLES

**Supplementary Table 1.** Examples of the calculation of repeat lengths.

| Sequence | Repeat length | Comment |
| --- | --- | --- |
| accaccac | 8 | The third iteration is only partial but still counts. |
| accac | 5 |  |
| acca | 2 | if followed by c then 4 |
| aaa | 3 |  |
| aca | 0 | if followed by c then 3 |
| acc | 2 | cc |
| tt | 2 |  |
| at | 0 |  |
| aatcagtt | 2+2 | aa + tt |

**Supplementary Table 2.**
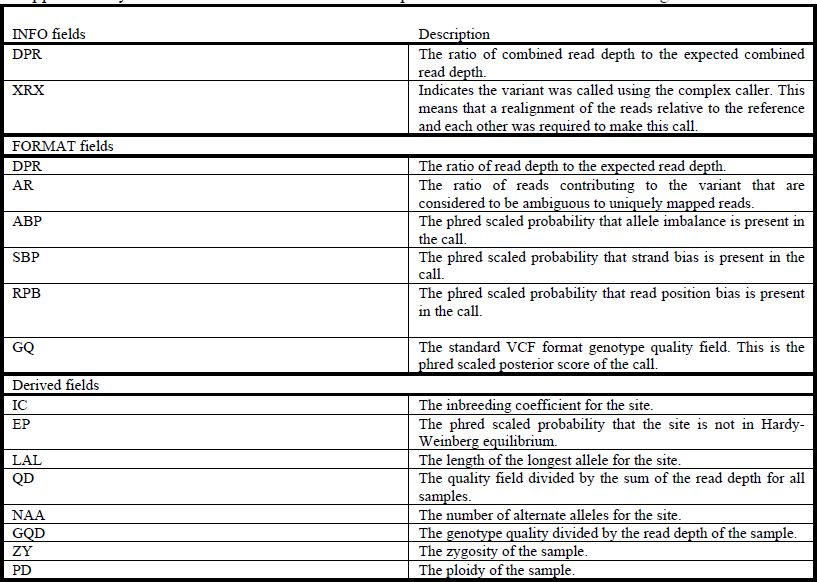
Attributes used in the Adaptive Variant Recalibration scoring model.

**Supplementary Table 3.** Conditional probability table for Mendelian inheritance of autosomal SNPs.

| $G_f$ | $G_m$ | $G_1$ | $P(G_1\vert G_f, G_m)$ |
| --- | --- | --- | --- |
| $\langle A, C \rangle$ | $\langle A, C \rangle$ | $\langle A, A \rangle$ | $1/4$ |
| | | $\langle A, C \rangle$ | $1/2$ |
| | | $\langle C, C \rangle$ | $1/4$ |

**Supplementary Table 4.** Examples of probabilities of genotypes matching the phasing framework by chance given a distribution of parent and offspring genotypes.

| Genotype Counts |  |  |  |  | Probability |
| --- | --- | --- | --- | --- | --- |
| 0/0 | 0/1 | 1/1 | 0/2 | 1/2 |  |
|  |  | 13 |  |  | 1 |
| | 13 | | | | $3.01 \times 10^{-1}$ |
| 6 | 7 | | | | $1.01 \times 10^{-2}$ |
| 1 | 12 | | | | $1.11 \times 10^{-1}$ |
| 1 | 11 | 1 | | | $1.36 \times 10^{-2}$ |
| 4 | 4 | 5 | | | $5.53 \times 10^{-4}$ |
| | 3 | 3 | 3 | 4 | $6.13 \times 10^{-5}$ |
| | 1 | 3 | 3 | 12 | $3.68 \times 10^{-1}$ |
| 1 | 5 | | 6 | 1 | $2.75 \times 10^{-4}$ |
| 1 | 11 | | 13 | 1 | $7.46 \times 10^{-2}$ |

**Supplementary Table 5.** Phase consistency of variants called on the CEPH 1463 large family with the GATK-UG v1.7.

| Call Set | Raw |  | 1<sup>st</sup> Tranche |  |
| --- | --- | --- | --- | --- |
|  | n | % | n | % |
| Phase consistent | 6,941,213 | 68.34 | 5,863,035 | 96.0 |
| Phase inconsistent | 2,263,975 | 22.29 | 184,169 | 3.01 |
| Repaired | 951,682 | 9.36 | 59,592 | 0.97 |
| Calls inside<br>phased segments | 10,156,870 | 99.53 | 6,106,796 | 99.98 |

**Supplementary Table 6.** Phase consistency of variants called on the CEPH 1463 large family provided by CGI.

| Call Set | Raw |  |
| --- | --- | --- |
|  | n | % |
| Phase consistent | 4,267,394 | 59.08 |
| Phase inconsistent | 1,384,465 | 19.16 |
| Repaired | 1,570,969 | 21.75 |
| Calls inside<br>phased segments | 7,222,828 | 99.98 |

**Supplementary Table 7.** *de novo* mutations called in offspring of 3^rd^ generation as a function of the number of siblings included in the calling.

| No. of siblings | All *de novos* (n) | SNV *de novos* (n) |
| --- | --- | --- |
| 0 | 1376 | 834 |
| 1 | 1129 | 719 |
| 2 | 1045 | 701 |
| 10 | 833 | 748 |

**Supplementary Table 8.** Variant statistics and metrics for CEPH 1463 pedigree called as a multigenerational pedigree.

| Sample | NA12877 | NA12878 | NA12879 | NA12880 | NA12881 | NA12882 | NA12883 | NA12884 | NA12885 | NA12886 | NA12887 | NA12888 | NA12889 | NA12890 | NA12891 | NA12892 | NA12893 |
| --- | --- | --- | --- | --- | --- | --- | --- | --- | --- | --- | --- | --- | --- | --- | --- | --- | --- |
| SNPs | 3,301,633 | 3,293,694 | 3,294,445 | 3,307,836 | 3,315,606 | 3,442,940 | 3,436,508 | 3,432,916 | 3,304,012 | 3,441,154 | 3,318,155 | 3,442,892 | 3,276,699 | 3,325,922 | 3,254,093 | 3,297,396 | 3,447,544 |
| MNPs | 32,935 | 32,297 | 32,262 | 32,178 | 32,242 | 32,007 | 31,518 | 31,652 | 31,827 | 31,536 | 32,157 | 31,948 | 31,980 | 31,471 | 31,464 | 31,760 | 31,906 |
| Insertions | 239,069 | 241,607 | 237,219 | 238,436 | 239,106 | 235,264 | 234,862 | 234,838 | 237,117 | 235,081 | 239,081 | 236,434 | 234,239 | 233,557 | 226,064 | 229,027 | 236,483 |
| Deletions | 237,252 | 239,207 | 234,799 | 235,378 | 235,886 | 233,941 | 233,308 | 233,327 | 235,753 | 233,762 | 237,162 | 234,291 | 229,733 | 237,391 | 227,668 | 231,186 | 234,930 |
| Indels | 79,615 | 78,645 | 78,439 | 78,812 | 78,893 | 77,568 | 77,331 | 76,981 | 78,630 | 76,970 | 78,745 | 77,230 | 72,713 | 72,690 | 70,970 | 73,147 | 77,694 |
| *de novos* | 162,77 | 11,656 | 5,227 | 5,943 | 7,107 | 170,438 | 167,930 | 168,946 | 4,752 | 170,791 | 5,886 | 170,476 | N/A | N/A | N/A | N/A | 172,377 |
| SNP |  |  |  |  |  |  |  |  |  |  |  |  |  |  |  |  |  |
| Ti/Tv | 2.11 | 2.11 | 2.11 | 2.11 | 2.11 | 1.97 | 1.97 | 1.97 | 2.11 | 1.97 | 2.11 | 1.97 | 2.11 | 2.11 | 2.12 | 2.12 | 1.97 |
| SNP |  |  |  |  |  |  |  |  |  |  |  |  |  |  |  |  |  |
| Het/Hom | 1.61 | 1.56 | 1.55 | 1.58 | 1.58 | 1.6 | 1.6 | 1.57 | 1.56 | 1.6 | 1.59 | 1.57 | 1.61 | 1.58 | 1.55 | 1.56 | 1.57 |

**Supplementary Table 9.** Variant metrics for WES at different coverage configurations and variant callers. FC, full coverage; RC, reduced coverage of parents; ARC, all reduced coverage.

| Family Caller (FC) | n | Ti/Tv | Het/Hom | % dbSNP |
| --- | --- | --- | --- | --- |
| 100X NA12891 | 40,444 | 2.59 | 1.55 | 92.30 |
| 100X NA12892 | 40,626 | 2.57 | 1.55 | 92.12 |
| 100X NA12878 | 40,927 | 2.6 | 1.55 | 92.53 |
| Family Caller (RC) | n | Ti/Tv | Het/Hom | % dbSNP |
| 50X NA12891 | 40,983 | 2.58 | 1.55 | 91.90 |
| 50X NA12892 | 41,346 | 2.57 | 1.54 | 91.76 |
| 100X NA12878 | 41,662 | 2.59 | 1.55 | 92.36 |
| Family Caller (ARC) | n | Ti/Tv | Het/Hom | % dbSNP |
| 50X NA12891 | 40,942 | 2.59 | 1.56 | 91.94 |
| 50X NA12892 | 41,289 | 2.57 | 1.54 | 91.85 |
| 50X NA12878 | 41,680 | 2.62 | 1.55 | 92.55 |
| Singleton (FC) | n | Ti/Tv | Het/Hom | % dbSNP |
| 100X NA12891 | 40,191 | 2.62 | 1.45 | 92.73 |
| 100X NA12892 | 40,455 | 2.61 | 1.47 | 92.59 |
| 100X NA12878 | 40,561 | 2.61 | 1.48 | 92.83 |
| Singleton (ARC) | n | Ti/Tv | Het/Hom | % dbSNP |
| 50X NA12891 | 39,689 | 2.63 | 1.41 | 92.45 |
| 50X NA12892 | 40,056 | 2.62 | 1.43 | 92.44 |
| 50X NA12878 | 40,202 | 2.64 | 1.45 | 92.92 |

**Supplementary Table 10.**
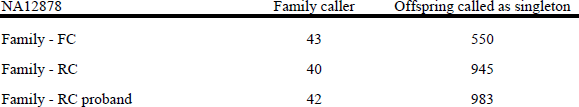
Mendelian error counts NA12878 WES in trio using different coverage configurations and callers.

| NA12878 | Family caller | Offspring called as singleton |
| --- | --- | --- |
| Family - FC | 43 | 550 |
| Family - RC | 40 | 945 |
| Family - RC proband | 42 | 983 |

**Supplementary Table 11.**
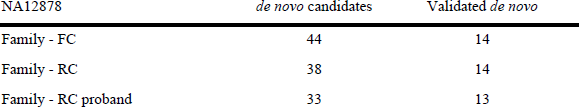
*de novo* mutations in NA12878 WES sample at different coverage configurations and callers.

## SUPPLEMENTARY FIGURES

**Supplementary Figure 1.**
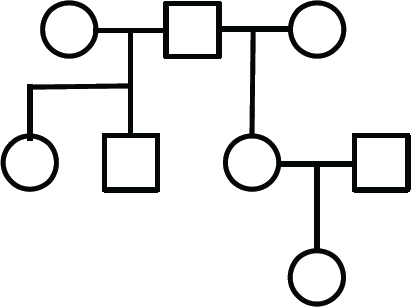
Complex three-generation pedigree including half-sibs.

**Supplementary Figure 2.**
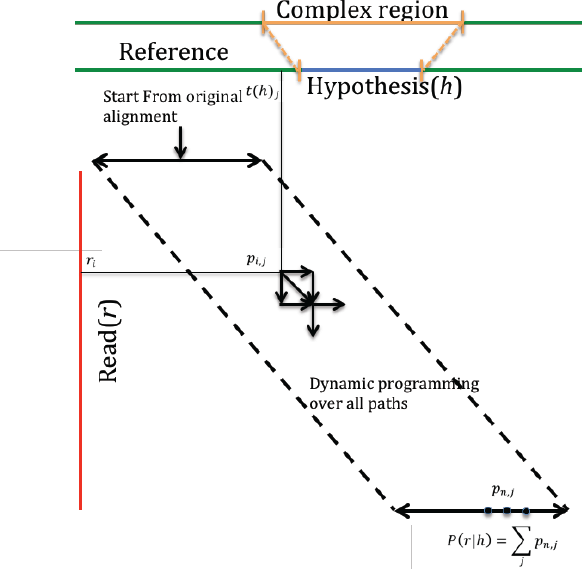
Alignment of all paths against template including an hypothesis.

**Supplementary Figure 3.**
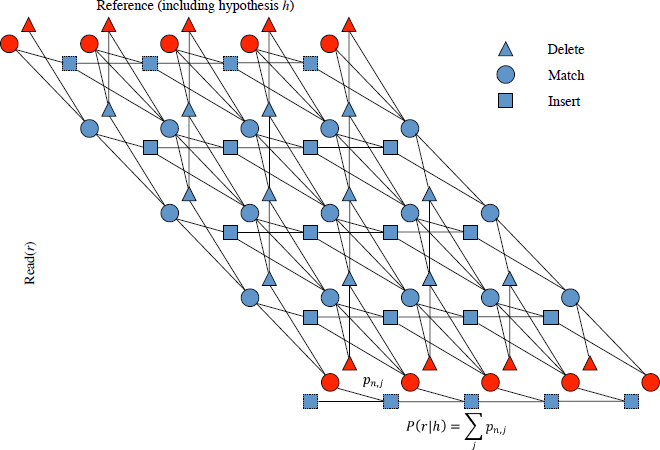
Dynamic programming for read alignments including insertion and deletion states.

**Supplementary Figure 4.**
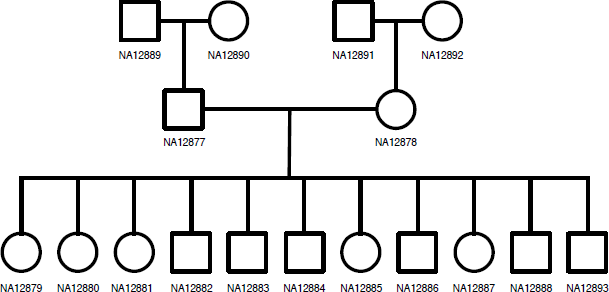
CEPH/Utah Pedigree 1463.

**Supplementary Figure 5.**
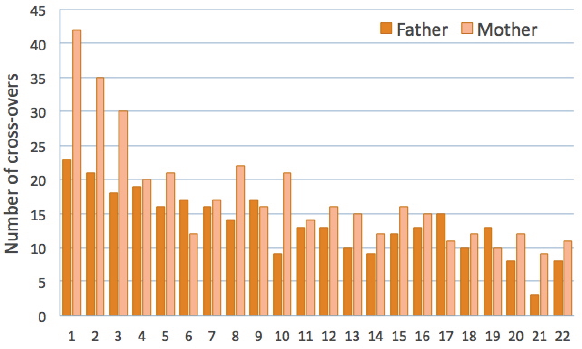
Recombination cross-overs identified across chromosomes of the CEPH 1463 large family offspring derived from each of their parents.

**Supplementary Figure 6.**
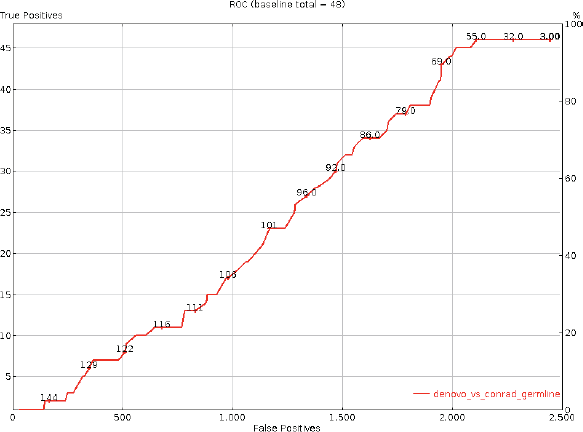
ROC curve of *de novo* mutations identified in the NA12878 sample vs validated *de novo* mutation baseline. DNP score shown along the curve.

**Supplementary Figure 7.**
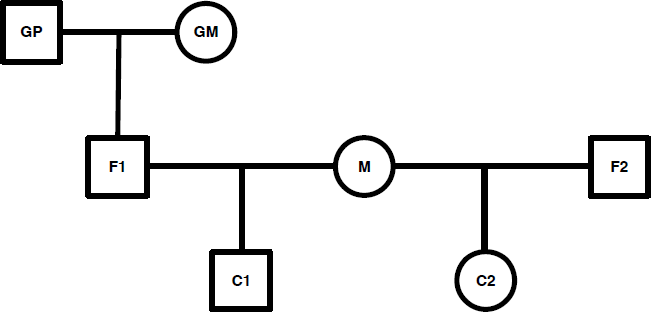
Extended pedigree structure used for simulations.

**Supplementary Figure 8.**
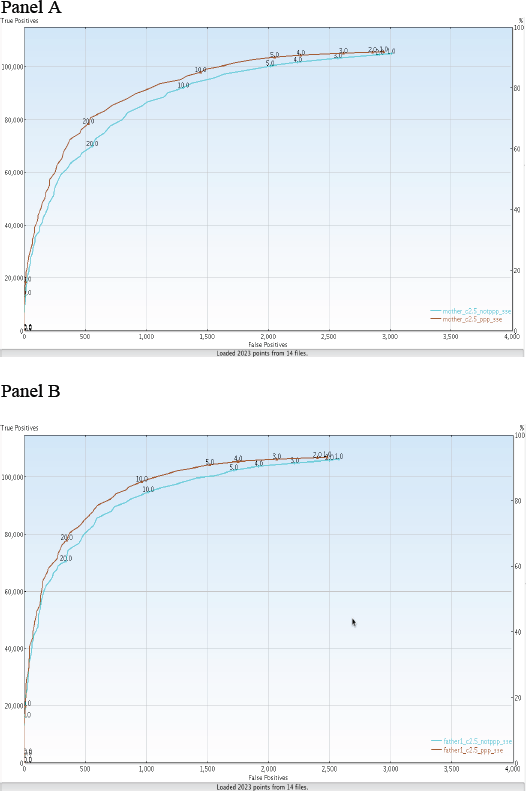
ROC curves for multigenerational pedigree simulations. A) ROC for mother (M) and B) ROC for Father 1 (F1) of pedigree in Supplementary Figure 7.

**Supplementary Figure 9.**
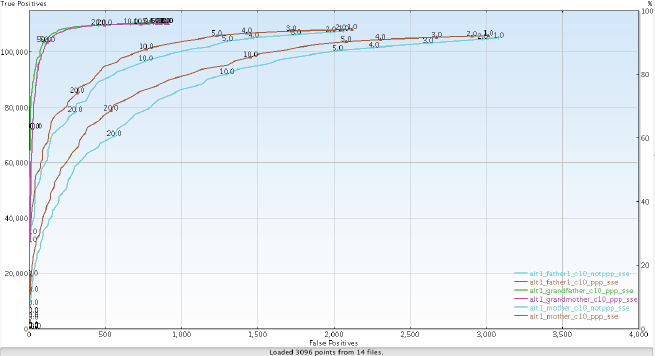
ROC curve for multigenerational pedigree simulation when grandparents (GP1 & GP2) depth is 30X and the rest of the pedigree is 5X.

**Supplementary Figure 10.**
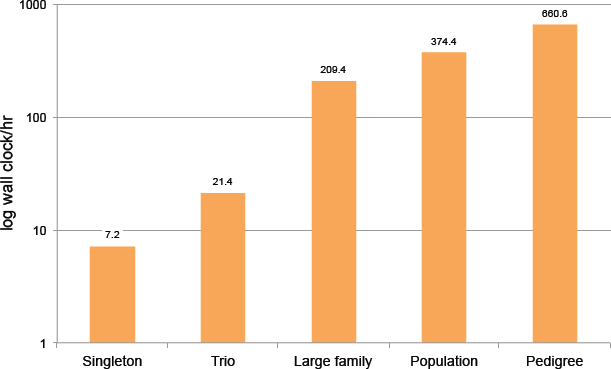
Cumulative wall times for different variant calling configurations on a commodity Linux server.

**Supplementary Figure 11.**
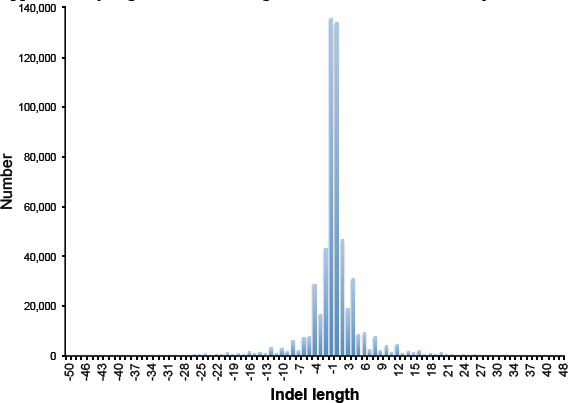
Indel length distribution in NA12878 joint trio calls.

## REFERENCES

Genomes Project Consortium et al. (2010) A map of human genome variation from population-scale sequencing. Nature 467, 1061–1073.

Genomes Project Consortium et al. (2012) An integrated map of genetic variation from 1,092 human genomes. Nature, 491, 56–65.

Ajay,S.S. et al. (2011) Accurate and comprehensive sequencing of personal genomes. Genome Res., 21, 1498–1505.

Ball,M.P. et al. (2012) A public resource facilitating clinical use of genomes. Proc. Natl. Acad. Sci. U.S.A.

Breiman,L. (2001) Random Forests. Machine Learning, 45, 5–32.

Brown,D.G. et al. (2004) A tutorial of recent developments in the seeding of local alignment. J Bioinform Comput Biol, 2, 819–842.

Cartwright,R.A. et al. (2012) A family-based probabilistic method for capturing de novo mutations from high-throughput short-read sequencing data. Stat Appl Genet Mol Biol, 11.

Conrad,D.F. et al. (2011) Variation in genome-wide mutation rates within and between human families. Nat. Genet., 43, 712–714.

DePristo,M.A. et al. (2011) A framework for variation discovery and genotyping using next-generation DNA sequencing data. Nat. Genet., 43, 491–498.

Drmanac,R. et al. (2009) Human Genome Sequencing Using Unchained Base Reads on Self-Assembling DNA Nanoarrays. Science, 327, 78–81.

Garrison,E. and Marth,G. (2012) Haplotype-based variant detection from short-read sequencing. arXiv.

Gilissen,C. et al. (2012) Disease gene identification strategies for exome sequencing. 1–8.

Gotoh,O. (1982) An improved algorithm for matching biological sequences. Journal of molecular biology, 162, 705–708.

Kidd,J.M. et al. (2008) Mapping and sequencing of structural variation from eight human genomes. Nature, 453, 56–64.

Koller,D. and Friedman,N. (2009) Probabilistic Graphical Models MIT Press.

Kong,A. et al. (2010) Fine-scale recombination rate differences between sexes, populations and individuals. Nature, 467, 1099–1103.

Levy,S. et al. (2007) The diploid genome sequence of an individual human. Plos Biol, 5, e254.

Li,B. et al. (2012) A Likelihood-Based Framework for Variant Calling and De Novo Mutation Detection in Families. PLoS Genet., 8, e1002944.

Li,H. et al. (2008) Mapping short DNA sequencing reads and calling variants using mapping quality scores. Genome Res.

Li,H. et al. (2009) The Sequence Alignment/Map format and SAMtools. Bioinformatics, 25, 2078–2079.

Manolio,T.A. et al. (2009) Finding the missing heritability of complex diseases. Nature, 461, 747–753.

Marth,G.T. et al. (1999) A general approach to single-nucleotide polymorphism discovery. Nat. Genet., 23, 452–456.

McCarroll,S.A. et al. (2008) Integrated detection and population-genetic analysis of SNPs and copy number variation. Nat. Genet., 40, 1166–1174.

McKernan,K.J. et al. (2009) Sequence and structural variation in a human genome uncovered by short-read, massively parallel ligation sequencing using two-base encoding. Genome Res., 19, 1527–1541.

ORawe,J. et al. (2013) Low concordance of multiple variant-calling pipelines: practical implications for exome and genome sequencing. Genome Medicine, 5, 28.

Pearl,J. (1988) Probabilistic Reasoning in Intelligent Systems Morgan Kaufmann.

Peng,G. et al. (2013) Rare variant detection using family-based sequencing analysis. Proc. Natl. Acad. Sci. U.S.A.

Ramu,A. et al. (2013) denovoGear:. Nat Meth, 1–5.

Saunders,C.J. et al. (2012) Rapid Whole-Genome Sequencing for Genetic Disease Diagnosis in Neonatal Intensive Care Units. Sci Transl Med, 4, 154ra135–154ra135.

Shafer,G.R. and Shenoy,P.P. (1990) Probability propagation. Ann Math Artif Intell, 2, 327–351.

Talkowski,M.E. et al. (2012) Clinical Diagnosis by Whole-Genome Sequencing of a Prenatal Sample. N Engl J Med, 367, 2226–2232.

Veltman,J.A. and Brunner,H.G. (2012) De novo mutations in human genetic disease. Nat. Rev. Genet., 13.

Zook,J.M. et al. (2013) Integrating sequencing datasets to form highly confident SNP and indel genotype calls for a whole human genome. arXiv.

